# Injury-induced electrochemical coupling triggers regenerative cell proliferation

**DOI:** 10.1101/2025.04.03.647033

**Authors:** J Liu, E Nerli, C Duclut, S A Vishen, N Berbee, S Kaufmann, C Ponce, B A Arrenberg, F Jülicher, R Mateus

## Abstract

Organ injury triggers non-neuronal electric currents essential for regeneration. Yet, the mechanisms by which electrical signals are generated, sensed and transmitted upon damage to promote organ growth remain unclear. Here, we uncover that organ regeneration relies on dynamic electrochemical coupling between tissue-wide depolarization and intracellular proliferative signalling. By sub-second live imaging of injured zebrafish larval fins, we identify events across timescales: a millisecond tissue depolarization gradient, followed by seconds-persistent intracellular Calcium responses. Subsequently within one hour, Voltage Sensing Phosphatase activity translates depolarisation into proliferation. Connecting these timescales with an electro-diffusive model showed that ionic fluxes and electric potential become coupled in the fin’s interstitial space, enabling organ-wide signal spreading. Our work reveals the coupling between fast electrical signals and slower intracellular signalling, ensuring organ regeneration.

## Introduction

Membrane potential (V_mem_) is a ubiquitous and essential property of all cells, established by ionic fluxes across membranes (*1*). While modulation of V_mem_ has been shown to have a direct impact in morphogenesis and regenerative processes (*2*–*12*), how these signals instruct specific cellular behaviours is still scarcely understood.

In epithelia, the differential distribution of ion pumps between the apical and basal cell membranes establishes a transepithelial potential (*13, 14*), also regulated by the presence of tight and gap junctions (*15*–*18*). Interestingly across fish species, perturbing ion flux invariably affects fin size (*19*–*27*), indicating a role for electrical signals in controlling processes such as cell proliferation and tissue growth. However, how V_mem_ changes can regulate proliferation during organ development and regeneration remains unknown. We therefore asked, how does organ damage affect tissue-scale bioelectrical states? How are electrical signals spatiotemporally generated in tissues and sensed by cells to promote organ (regenerative) growth?

Here, we applied UV-laser microdissection to precisely injure zebrafish larval fins, while live imaging this process with sub-second resolution. This allowed us to quantitatively map the spatiotemporal dynamics of V_mem_ and Calcium signals *in vivo*. We found that organ injury leads to an immediate tissue-scale membrane depolarization gradient, followed by a diffusive Calcium wave. To understand the onset and propagation of these signals at milli- to seconds timescales in the fin, we developed electro-diffusive theory, coupling ion fluxes to tissue-scale voltage and Calcium dynamics. This clarified that the observed signal spreading and persistence is rooted in ionic diffusion within the fin’s interstitial fluid. Manipulating V_mem_ and interstitial fluid proved sufficient to alter the spatiotemporal dynamics of the injury-induced Calcium response. Surprisingly, we found that membrane depolarisation triggers cell proliferation within 1h. At the molecular level, we find that signal transduction occurs due to a transmembrane protein, Voltage Sensing Phosphatase, which is necessary and sufficient to sense tissue depolarisation and trigger proliferation, enabling successful organ regeneration. Our findings uncover a bioelectric injury-sensing mechanism necessary to fast-track organ regeneration.

## Results

### A tissue-scale depolarization gradient precedes a calcium wavefront upon organ injury

In order to investigate the spatiotemporal dynamics of electrical signals elicited by organ damage, we established an *in vivo* live-imaging assay joining optical electrophysiology tools with precise UV-laser microdissection. This allowed milliseconds resolution of tissue-scale signal dynamics induced by injury, in the 2 days-post-fertilisation (dpf) zebrafish caudal fin (Methods).

The 2 dpf fin exhibits a planar tissue architecture, consisting of two adjacent epithelial layers that enclose fibroblast-like cells and structural elements within its interstitial fluid (Fig. 1A, Fig. S1C). The presence of tight junctions in the apical epithelial layer of the fin makes this tissue an effective barrier to the extraembryonic environment (*16*), guaranteeing high electrical resistance (*28*)(Fig. 1A’, Fig. S1A). Notably, we find that gap junctions are also restricted to this fin layer (Fig. 1A’,B; Fig. S1B), providing electrical coupling (*29*–*31*).

**Figure 1.**
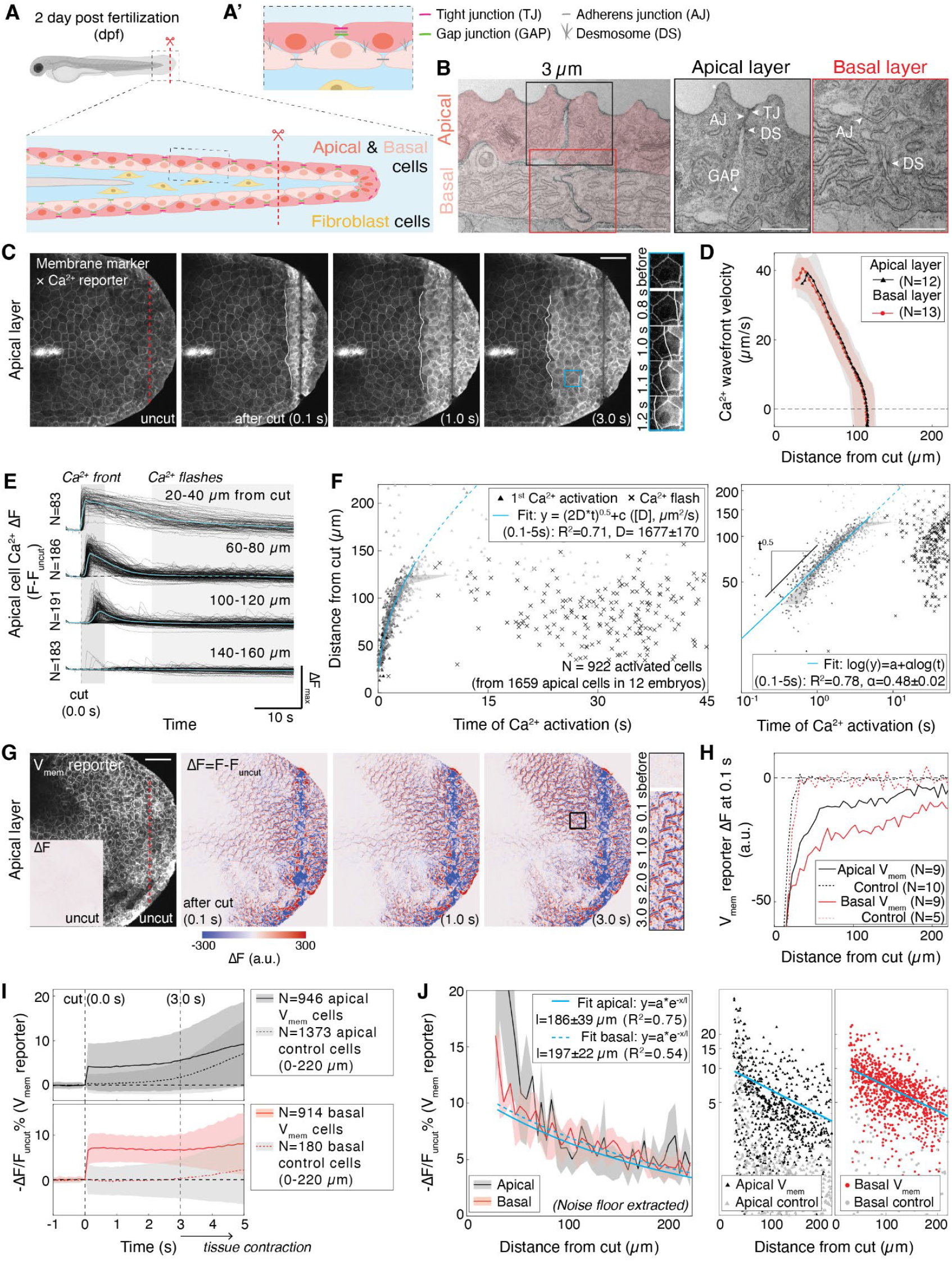
A membrane depolarization gradient and diffusive Ca^2+^ wavefront are rapid fin-wide injury responses. **(A)** Zebrafish larva at 2 days-post-fertilization (dpf), highlighting caudal fin architecture, junctions **(A’)**, and amputated region (red dash). **(B)** Electron transmission micrographs of 2dpf fin with annotated junctions in apical (black) and basal (red) epithelial layers. **(C)** 2dpf Fin before and after cut (red dash), in transgenics labeling intracellular Ca^2+^ (ubb:GCaMP6f) and membranes of apical epithelium (claudinb:Lyn-GFP). Blue inset, cellular detail. White lines, activated Ca^2+^ front. **(D)** Velocity of Ca^2+^ front vs. position relative to wound in apical (black) and basal (red) epithelia. Shades, SD. **(E)** Change of cytosolic Ca^2+^ intensity (ΔF, vs uncut) for individual cells vs. time. Cyan, average. **(F)** Position of apical epithelial cells relative to wound vs. time of Ca^2+^ wavefront (triangle) or flash (cross) activation. Right, log-to-log scale. Shade, 25%-75% percentile. Cyan lines, diffusive fit in linear scale (D: diffusion coefficient), linear fit (ɑ: scaling exponent) in log-to-log scale (0.1s-5s). Cyan dash, diffusive fit continuation (0.1s - 20s). **(G)** Left, 2dpf fin apical epithelium labelled with Voltron2 (V_mem_ reporter) before cut (red dash). Inset, change of Voltron2 intensity in uncut fins (ΔF, vs previous 0.1s uncut). Middle to right, change of Voltron2 intensity at 0.1s, 1.0s and 3.0s post-cut (ΔF, vs uncut). Black inset, cellular detail. **(H)** At 0.1s post-cut, changes of Voltron2 intensity (ΔF, vs uncut) vs. position relative to wound, for apical (black) or basal (red) epithelium, plus membrane reporters (control, dash). **(I)** Time traces of fractional drop of Voltron2 intensity before and after cut (-ΔF/F_0_ ; F_0_ = uncut intensity, depolarization readout) for apical (top) or basal (bottom) cells, plus membrane reporters (control, dash). Shades, SD. After ⩾3s pos-tinjury, Voltron2 cannot be rigorously detected due to tissue contraction (Methods). **(J)** At 0.1s post-cut, fractional drops of Voltron2 intensity (-ΔF/F_0_) in apical or basal cells vs. position relative to wound (same dataset as I). Left, Mean ±SD (shade), with noise floors (from membrane controls) extracted. Right, Voltron2 intensity in apical or basal cells, vs. membrane controls (gray). Cyan lines, exponential fits in linear (left) and log-to-linear (right) scales. Error, 95% CI of parameter fits. R^2^, goodness of fit. N: number of larvae (D,H) or cells (E,F,I,J). Scale bars: 50µm (C,G); 2µm, 1µm (B).

We first focused on quantifying the post-injury dynamics of intracellular Calcium, given previous reports that it composes the earliest wound signal that can direct cellular responses (*12, 32*–*38*). Applying our developed live imaging assay, we detected the formation of a tissue-scale Ca^2+^ wavefront as early as 100ms post-injury, initiated by wound-adjacent cells (Fig. 1C, Fig. S2A, Movie S1). Combining Ca^2+^ signalling transgenics with epithelial membrane reporter lines (Methods), we found this wavefront concomitantly propagates in both epithelial fin layers up to ∼120µm from the wound (apical: 118.4±14.3 µm, basal: 118.7±8.5 µm), slowing down and stopping at ∼5s after injury (apical: 4.9±0.9 s, basal: 5.3±1.0 s) (Fig. 1D, Movie S1). At 15s after injury, only wound-adjacent cells showed elevated Calcium levels. In the subsequent 30s, Ca^2+^ flashes occur in single epithelial cells traversed by the wavefront (Movie S1).

To determine the speed of the Ca^2+^ wavefront activation, we measured the total intracellular Calcium intensity change per cell (Fig. 1E, Fig. S2B), and extracted a characteristic time of Ca^2+^ cell activation (Methods). We found a robust power-law dependence between distance from wound and Ca^2+^ activation time, with a ∼0.5 exponent in both fin epithelial layers (apical cells: 0.48±0.02, basal cells: 0.46±0.01, Fig. 1F, Fig. S2C-D), indicating diffusive propagation (*39*) of the Ca^2+^ activation signal from the injury. We then extracted diffusion coefficients per layer (apical: 1677±170 µm^2^/s, basal: 1877±179 µm^2^/s, Fig. 1F, Fig. S2C-D), which remarkably are in comparable ranges to ion diffusivity in aqueous medium (Na^+^: 1334 µm^2^/s, K^+^: 1957 µm^2^/s, Cl^-^: 2032 µm^2^/s (*40*)). Notably, this diffusive propagation is invariant to differences in fin injury shape, size (Fig. S3A-B, Movie S2), or type (laser or mechanical cut; Fig. S3C, Movie S3), as long as the lesion punctures both fin epithelial layers. Thus, the observed Ca^2+^ dynamics point towards a physical mechanism that is rooted in tissue-scale ion fluxes altered by organ injury.

The sub-second Ca^2+^ wavefront dynamics occurring in both fin epithelial layers suggested the presence of a signal that could act faster than known chemical regulators upon fin injury (*41*). Motivated by the known coupling between Ca^2+^ signalling and voltage changes in neurons (*42, 43*), we asked whether the injury affects V_mem_. To probe changes in V_mem_ per fin layer, we expressed a FRET-based voltage reporter, Voltron2 (*44*), into apical or basal epithelial reporter transgenics (Fig. 1G, Fig. S4A, Methods) and applied our live-imaging injury assay.

At 100ms after injury, we detected widespread cell depolarisation in both fin layers, corresponding to a spatially-graded drop of Voltron2 fluorescence spanning ∼200µm from the wound (Fig. 1H; Fig. S4A-B). In contrast, at 100ms, intracellular Ca^2+^ is only activated within 20µm from the wound (Fig. 1C). Over the next 3s, Voltron2 intensity is maintained (Fig. 1I), suggesting that cells remain depolarized as the Ca^2+^ wavefront propagates. By quantifying the fractional change of Voltron2 intensity occurring before and after cut (ΔF/F_0_, Methods), we found that the spatial gradient of V_mem_ depolarization displays an exponential profile with comparable characteristic lengths *l* for both epithelial layers (apical: 186±39 µm, basal: 197±22 µm, Fig. 1J, Fig. S4C-D). These results establish that a tissue-scale depolarization gradient spatiotemporally precedes the Ca^2+^ wave in injured fins, and together, these multi-scale signals compose the earliest tissue-wide injury responses yet detected.

### Multicompartment electro-diffusion model explains voltage and Ca^2+^ injury responses

Our measurements of the fin injury responses led to two central questions: how does a local lesion induce a widespread (>100 µm) voltage response, within milliseconds (100 ms)? How do Ca^2+^ injury responses emerge on a different timescale than voltage, yet displaying activation dynamics similar to ionic diffusion? To address these, we developed electro-diffusion theory in multiple compartments, to solve for the changes of ion fluxes and electric potentials upon injury in the 2dpf caudal fin. Our theory minimally describes this organ as an electrically-coupled apical epithelial sheet that separates the high-osmolarity interstitial fluid (IF)(*45, 46*) from the low-osmolarity extraembryonic environment (Fig. 2A). Importantly, the presence of tight junctions (Fig. 1B, Fig. S1A), due to their inherent leakiness at this developmental stage (*16*), allows for paracellular ion fluxes between the interstitial and extraembryonic spaces (Fig. 2A, Sec. I of Supplementary Notes).

**Figure 2.**
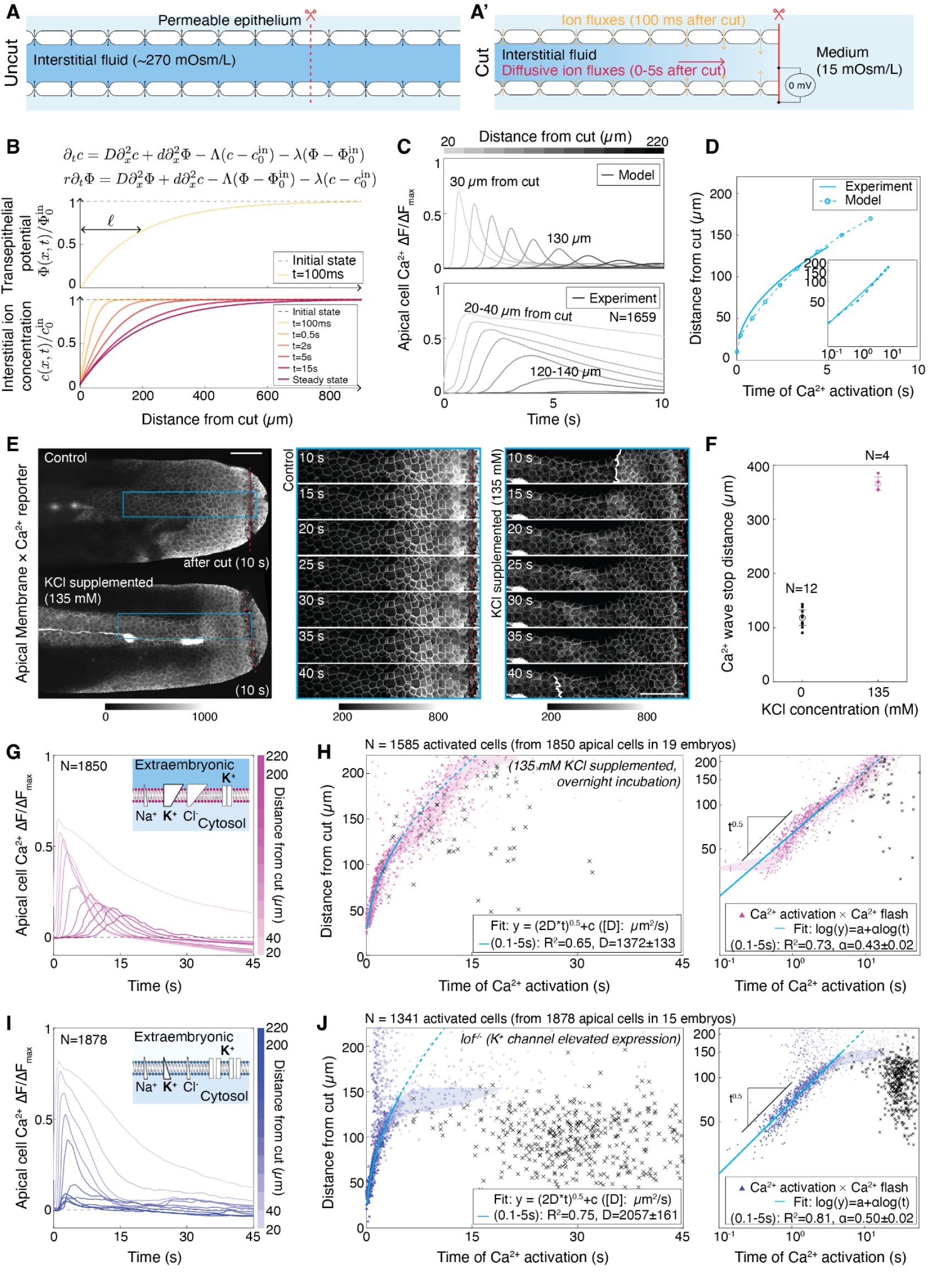
Potassium perturbations modulate tissue-scale Calcium dynamics upon injury. **(A)** Electro-diffusive model for the apical fin epithelium enclosing interstitial fluid (IF), contacting external medium, before and after cut **(A’). (B)** Top, predicted changes of transepithelial potential vs. position relative to wound within 100ms. Bottom, predicted changes of interstitial ionic concentration vs. position relative to wound between 100ms - 15s, with steady state profile. Model equations obtained using electroneutral approximation of Poisson-Nernst equations for two monovalent ionic species (see SI). *c*, IF ion concentration; *Φ*, electric potential energy. *c*_*0*_^*in*^ and *Φ*_*0*_^*in*^, respective values prior to wounding. *D* and *d*, average and half difference of the ion diffusivities, respectively. *Λ* and *λ*, average and half difference of epithelium permeability to each ion, times the volume-to-area ratio of the fin, respectively. *r* ≃ *10*^−*6*^ is the dimensionless timescale associated with electric potential relaxation. **(C)** Top, calculated changes of the cytosolic Ca^2+^ concentration vs. time from the model. Bottom, measurements of position-averaged changes of cytosolic Ca^2+^ intensity (ΔF/ΔF_max_, vs uncut, normalized per embryo) vs. time. **(D)** Diffusive fit to Ca^2+^ front position vs. time, in experiments (line, from Fig. 1F) vs model (dash). **(E)** 2dpf fins at 10s post-cut (red dash), in control (top) or KCl-treated (bottom) transgenics, ubb:GCamp6f (Ca^2+^ reporter), claudinb:Lyn-GFP (membrane marker). Right, Time-lapse from respective blue insets. White lines, Ca^2+^ front position. **(F)** Ca^2+^ front traverse distance in control vs. KCl-treated larvae. Error bars, SD. **(G, I)** Position-averaged changes of cytosolic Ca^2+^ intensity (ΔF/ΔF_max_, vs uncut, normalized per embryo) from apical fin epithelium vs. time, in KCl-treatment **(G)** or *lof*^*-/-*^ **(I)**. Insets: condition-specific depiction of ionic concentration (blue) and transport across compartments. **(H, J)** Position of apical epithelial cells relative to wound vs. time of Ca^2+^ wavefront (triangle) or flashes (cross) activation, in KCl-treated **(H)** or *lof*^*-/-*^ **(J)**. Shades, 25%-75% percentile. Cyan lines: diffusive fit in linear scale; linear fit in log-to-log scale (0.1s - 5s). Cyan dash, diffusive fit continuation (0.1s - 20s). Error, 95% CI of parameter fits. R^2^, goodness of fit. N: number of larvae (F) or cells (C,G,I). Scale bars: 100µm.

As the injury induces local short-circuit of the fin’s transepithelial potential, a sudden change occurs at the wound boundaries: the high and low osmolarity media from inside and outside the zebrafish become connected. This triggers local changes of V_mem_ that are spatiotemporally-coupled via electro-diffusive dynamics of cation (Na^+^) and anion (Cl^-^) species inside the fin (Fig. 2B, Sec. I of Supplementary Notes). Interestingly, our model shows that, upon injury, the electric potential and ion concentration change at different timescales.

The first response of the electric potential, occurring at milliseconds, is faster than ion diffusion (*40*). Hence, the dynamics of electric potential reduces to that of cable theory (*47, 48*). Here, cable theory represents the epithelium as a capacitor and resistor in parallel, taking into account the conductivity of the IF (Fig. 2B, Sec. II of Supplementary Notes). Upon injury, which corresponds to a short-circuit of the epithelium, the electric potential in the interstitial fluid relaxes to a quasi-steady state over 100 ms. After relaxation, it assumes an exponential profile that decays over a characteristic length 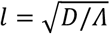, set by the diffusion constant of ions *D*, and the parameter *Λ*, describing leakage through the epithelium. By explicitly considering ion transport across the basolateral cell membrane facing the IF, we show that the potential difference across this membrane is proportional to the transepithelial potential, and therefore also assumes an exponential profile (Fig. 2B, top). Using diffusion constants of Na^+^ and Cl^-^, geometric parameters from the fin architecture (Fig. S1C-E), and the determined characteristic length *l* (Fig. 1J), we estimate *Λ* = *0*.*05 s*^−*1*^ (Sec. III of Supplementary Notes). For typical concentration of ions in the IF (*45, 46*) this corresponds to an epithelium resistance in unit area of ∼*200 Ωcm*^*2*^, well within the range of values reported for epithelial resistance (*28*).

At a slower timescale, seconds subsequent to injury, ion diffusion takes place from the high-concentration IF to low-concentration extraembryonic medium. Here, the dynamics of ions and electric potential become coupled, being well-described by effective diffusion equations with a source term stemming from ion transport across the epithelium (Fig. 2B, Sec. II of Supplementary Notes). As a result, an ionic concentration profile spreads along the fin’s anterior-posterior axis (Fig. 2B, bottom). This profile drives the intracellular Calcium dynamics, leading to the observed diffusive activation wave.

We also developed a minimal non-linear model for single cell Calcium responses that qualitatively captures the rising time and amplitude decrease (Fig. 2C, Sec. III of Supplementary Notes). This recapitulates an activation front of intracellular Calcium with a close-to-ion diffusion coefficient (Fig. 2D, Movie S4). Moreover, the dispersion of the experimental Calcium activation data (Fig. 1F, Fig. S2C) can be accounted for in a stochastic version of the model (Fig. S16, Sec. III of Supplementary Notes). Therefore, our model strongly suggests that the observed dynamics of Calcium signalling is driven by the electrical and ionic changes that take place inside the fin’s interstitial space. This is consistent with the observation that the Calcium wave occurs independently of extraembryonic Calcium in the media (Fig. S5A, Movie S5). Furthermore, the Calcium wavefront remains unaffected even when gap junction pore formation is blocked (Fig. S5B-E, Movies S6-7).

### Potassium flux remodels organ-scale injury responses

Our physical model suggests that the observed Calcium wavefront is rooted in the ionic changes occurring in the fin upon injury. We thus asked whether changing ion composition and fluxes would spatiotemporally affect this injury response. We focused on potassium (K^+^), since this ion is both central to determining V_mem_ in cells (*49*), and an important regulator of fish fin sizes through specific leak channels (*20, 25*). Considering this, we incorporated K^+^ into our model (Sec. IV of Supplementary Notes), predicting that K^+^ perturbations would lead to two prominent tissue-level changes: i) IF ionic composition and conductivity, and ii) V_mem_.

In the first case, given the fin’s epithelial permeability (Fig. 2A)(*16*) and its low IF K^+^ concentration (*45, 46*), then high extraembryonic K^+^ concentration should change the IF cation composition. This would result in an altered electro-diffusive Ca^2+^ response upon injury (Sec. IV of Supplementary Notes). To verify this, we incubated larvae in highly concentrated potassium chloride (KCl) (Methods). Remarkably, by inducing fin injury in this condition, the Calcium wavefront propagated for more than 300µm, 3-fold longer than wildtype (Fig. 2E-H, Movies S8-9). This Ca^2+^ wavefront also propagated with a smaller diffusion coefficient compared to wildtype (Fig. 2H), consistent with the theory (Sec. IV of Supplementary Notes).

Second, our theory also predicts a direct effect of K^+^ supplement on V_mem_ (Sec. IV of Supplementary Notes), with experimental evidence indicating that it would lead to cell depolarization (*50*–*52*). Therefore, we investigated whether high extraembryonic KCl affects Ca^2+^ flashes, as reported in neuronal systems (*43, 53*). Interestingly, we observed a tissue-scale reduction of wound-induced Ca^2+^ flashes (Fig. 2H, Fig. S6B, S7C, Movie S9). Conversely, enhancing K^+^ efflux in injured fins with two known functional assays (*longfin*^*t2*^ homozygous mutants, i.e. *lof*^-/-^ (*25*) or FK506 treatment (*21, 23, 54*)) led to a prominent Ca^2+^ flash increase (Fig. 2J, Fig. S6C-D, S7D, Movie S9). The Ca^2+^ activation wavefront, on the other hand, remained mostly unaffected in these conditions (Fig. 2I-J, Fig. S6C-D, S7A-B, compared to Fig. 1F). Taken together, we conclude that K^+^ flux modulation is sufficient to drastically remodel Ca^2+^ injury responses, by regulating IF ionic composition and V_mem_.

### Membrane potential regulates proliferation and tissue regeneration

Our results show that electrical signals in the form of tissue-wide depolarisation arise upon organ injury. However, organ regeneration relies on cell proliferation to recover functionality, occurring over hours or even days (*55, 56*). While electrical cues have been shown to trigger diverse cell behaviours in injured tissues (*12*), it remains unclear how cells react to fast changes of electrical signals to control slower organ regenerative growth. To fill this gap, we extended our experimental assay to explore how spatial V_mem_ changes triggered by injury could be linked to proliferation and regeneration dynamics occurring in the 2dpf caudal fin (Fig. 3A).

**Figure 3.**
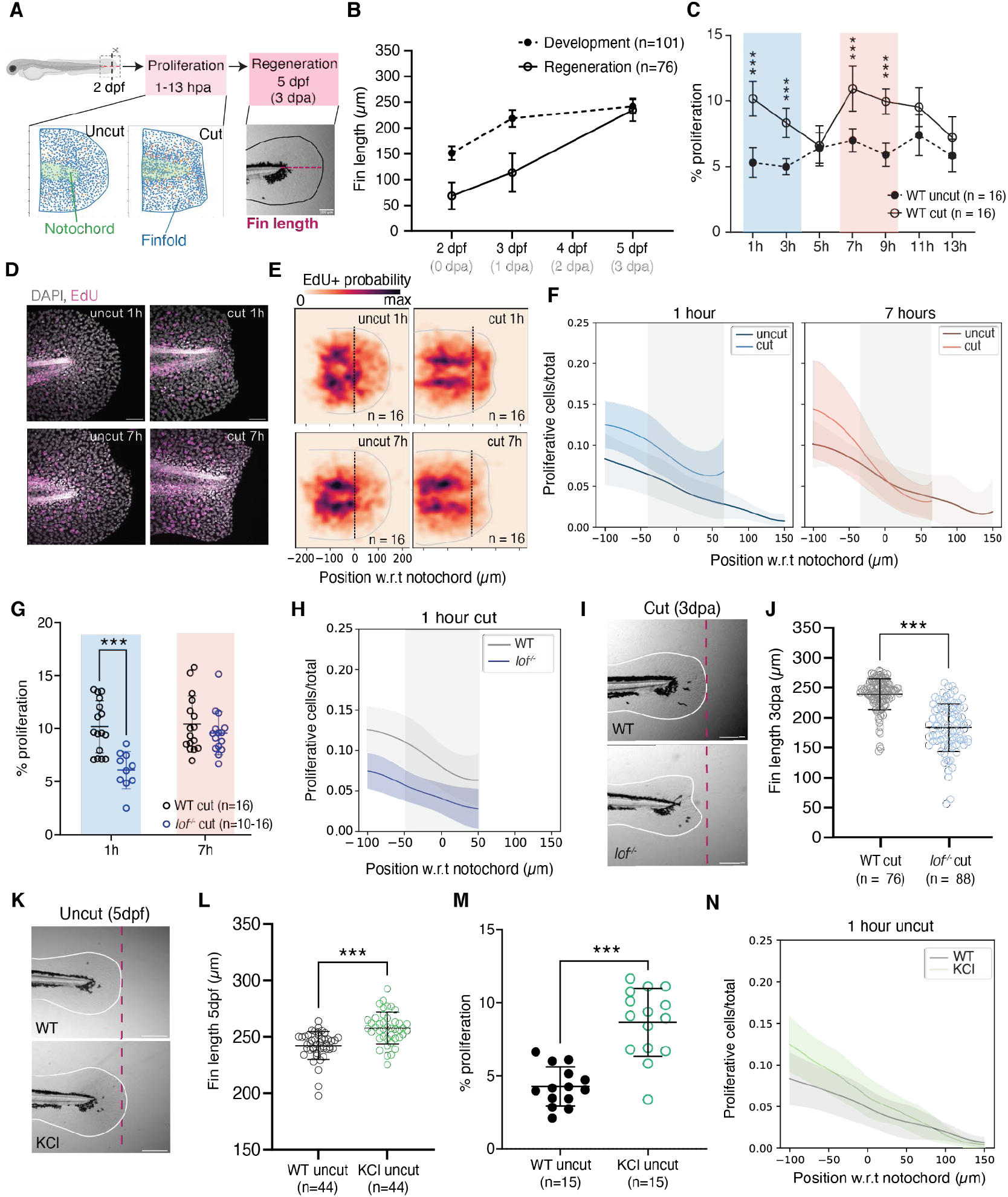
V_mem_ changes regulate cell proliferation at 1-hour post-injury. **(A)** Proliferation and regeneration assays. **(B)** Fin lengths in development vs regeneration, 2-5 dpf. **(C)** Percentage of proliferative cells (EdU+) per total (DAPI+), in uncut and cut fins from D. Shades, proliferation peaks. Mean ±95% CI. **(D)** Immunostainings labeling proliferative cells (magenta, EdU+) and nuclei (grey, DAPI). **(E)** Heatmaps displaying probability of occurrence of a proliferative cell within fin area. Outline, average fin boundary. Dash, notochord tip. **(F)** Fraction of proliferative cells vs. position, in cut vs uncut fins, at 1h and 7h. Note increase within 100µm from cut (grey rectangle) in 1 hpa vs. 7 hpa. **(G)** Percentage of proliferative cells per total in *lof*^-/-^ vs. WT, at 1hpa and 7hpa. Mean ±95% CI. **(H)** Fraction of proliferative cells vs. position, in WT vs *lof*^-/-^, at 1 hpa. **(I)** WT and *lof*^-/-^ regenerated fins. **(J)** Fin lengths in regenerated WT vs. *lof*^-/-^. **(K)** Uncut fins in WT and KCl-treated larvae. **(L)** Fin lengths in uncut WT vs. KCl-treated larvae. **(M)** Percentage of proliferative cells per total in uncut WT vs. KCl-treated larvae. **(N)** Fraction of proliferative cells vs. position, in uncut WT and KCl-treated larvae. Scale bars: 50µm (C,G,M); 200µm (I,K). I,K: pink dash, 5dpf uncut WT fin length. F,H,N: line and shade, mean ±SD; zero, notochord tip. B,J,L,M: mean ±SD. All statistics: ***p≤0.001; C,G,J: two-tailed, unpaired, non-parametric Mann-Whitney tests; L: two-tailed, unpaired, parametric t-test. n, number of embryos.

After injury at 2dpf, the zebrafish caudal fin regrows within 3 days (Fig. 3B, Fig. S8A-D)(*55, 57*), with proliferation reported to occur mostly within the first day of regeneration (*55*). Surprisingly, by labeling proliferative cells (EdU+), we found two proliferation peaks occurring within 13h post-injury: the first at 1-3 hours-post-amputation (hpa) and the second at 7-9 hpa (Fig. 3C). Interestingly, spatial analysis showed that between 1-3 hpa, proliferation is higher within ∼100µm from the wound, when compared to 7-9 hpa (Fig. 3D-F, Fig. S8E-F).

The similar tissue length scales between proliferation and graded depolarization made us ask whether electrical signals arising upon tissue injury (Fig. 1) could trigger proliferation in a spatiotemporal dependent manner. To test this, we measured proliferation as well as regenerative ability (Fig. 3A) under the ion flux perturbations tested previously (Fig. 2). Remarkably, in *lof*^*-/-*^ and upon FK506-treatment, the first proliferative peak at 1 hpa was impaired, without affecting proliferation at 7 hpa (Fig. 3G-H, Figs. S9F, S10F-G). This resulted in fin length defects in the first day post amputation (Figs. S9B-C’, S10A-C’), ultimately leading to incomplete regeneration at 3 dpa (Fig. 3I-J; Figs. S9B,D-E, S10B,D-E). These phenotypes are regeneration-specific, since these perturbations did not influence developmental proliferation or fin length (Figs. S9A-B, S10A-B,H). Hence, proliferation occurring in the first hour post-injury is affected by changes in electrical signals, necessary for successful regeneration.

To understand whether V_mem_ depolarisation can trigger organ growth through increased proliferation, we induced tissue-wide depolarisation by KCl-supplemented medium for 1h (*50*– *52*), in uncut and cut fins (Methods). Surprisingly, in the absence of injury, this one hour treatment led to fin overgrowth (Fig. 3K-L, 10% length increase) by amplifying fin-wide proliferation (Fig. 3M-N), without affecting body size (Fig. S11E). Longer KCl treatment (Methods) also led to longer fins (Fig. S11F). Under injury conditions, KCl treatment still resulted in successful fin regeneration (Fig. S11A-D). Overall, our results indicate that electric signals modulate fin growth and regeneration, and remarkably, tissue depolarisation is sufficient to induce proliferation independently of wounds.

### Voltage Sensing Phosphatase relays V_mem_ changes into fin proliferation

Next, we set out to investigate the molecular mechanism that could sense depolarisation, transmitting such signals intracellularly to affect the first proliferation peak. Previous *in vitro* studies suggested that changes in V_mem_ can trigger proliferation through membrane lipids remodelling (*58*), but the molecular effector remains unknown. Therefore, we probed the literature for candidate membrane proteins with voltage sensing domains, coming across the protein Voltage Sensing Phosphatase (VSP)(*59*), encoded in zebrafish by *tpte* (*60, 61*). This protein functions both as a voltage sensor and phosphatase, having as substrates the lipids phosphatidylinositol (3,4,5)-trisphosphate (PIP_3_) and phosphatidylinositol 4,5-bisphosphate (PIP_2_). Importantly, VSP changes conformation upon membrane depolarisation above ∼40 mV, activating its cytosolic phosphatase, dephosphorylating PIP_3_ and PIP_2_ (*59, 62, 63*). Consequently, we hypothesized that VSP senses and transduces injury depolarisation signals, leading to the observed fin proliferation peak at 1 hour post amputation (Fig. 3C,F).

Interestingly at 2dpf, we found that VSP is present in the caudal fin, being exclusively localized at the membranes of apical epithelial cells (Fig. 4A). This prompted us to further investigate whether proliferation is spatially biased across the two epithelial fin layers. While in development, proliferation occurs mostly in the basal epithelium (*64*), surprisingly, during regeneration, we found that the fin’s apical epithelial layer is the main tissue proliferating (Fig. 4B).

**Figure 4.**
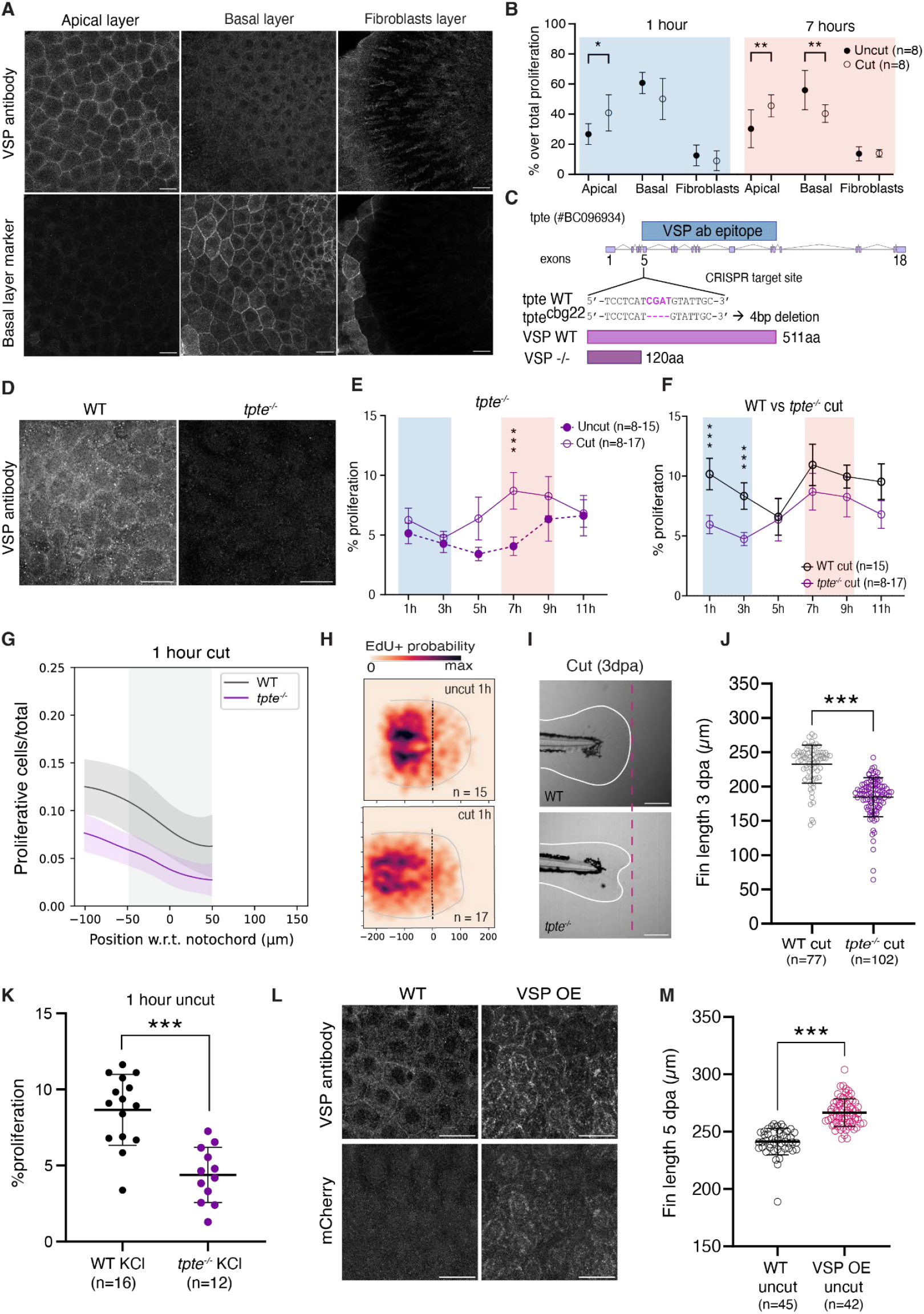
VSP translates changes in V_mem_ into proliferation. **(A)** VSP immunostainings in 2dpf apical and basal fin epithelial layers (single frames). Basal layer marker, tp63:CAAX-GFP. **(B)** Percentage of proliferating cells in each layer per total fin proliferation, in uncut and cut conditions. Mean ±95% CI. **(C)** *tpte* genomic structure. Target sequence of *tpte*^*cbg22*^ mutant in exon 5, with truncated VSP protein. VSP antibody epitope is indicated. **(D)** VSP immunostainings in 2dpf WT and *tpte*^*-/-*^ fins (max intensity projections). **(E-F)** Proliferation percentages over total cells between 1-11 hpa, in *tpte*^*-/-*^ uncut vs. cut **(E)** or in WT vs. *tpte*^*-/-*^ **(F)**. Mean ±95% CI. Shades, WT proliferation peaks from Fig. 3C. **(G)** Fraction of proliferative cells vs. position in WT vs *tpte*^-/-^, at 1hpa. Zero, notochord tip. Grey area, 100µm from cut. Line and shade: mean ±SD. **(H)** Heatmaps displaying probability of occurrence of a proliferative cell within fin area, in 1h cut and uncut *tpte*^*-/-*^. Outline, average fin boundary. Dash, notochord tip. **(I)** WT and *tpte*^*-/-*^ cut fins. Outlines, fin boundary. Pink dash, 5dpf uncut WT fin length. **(J)** Fin lengths in WT vs. *tpte*^*-/-*^. **(K)** Percentage of proliferative cells per total in *tpte*^*-/-*^ vs. WT, KCl-treated. Mean ±95% CI. **(L)** VSP immunostaining in VSP overexpression transgenics (VSP-OE, mCherry) and WT, 2dpf apical epithelium. MIP of two planes. **(M)** Fin lengths from WT vs. VSP-OE. Scale bars: 20µm (A, D, L); 200µm (I). n, number of embryos. J,K,M: mean ±SD. All statistics: *p≤0.05, **p≤0.01, ***p≤0.001; B,E,F,K: two-tailed, unpaired, non-parametric Mann-Whitney tests; J,M: two-tailed, unpaired, parametric t-tests.

To directly test whether VSP is responsible for the fin proliferative response upon injury, we generated a *tpte* CRISPR mutant that carries a stop codon in the 5’ region of its coding sequence (*tpte*^*cbg22*^, Fig. 4C), leading to the absence of VSP expression (Fig. 4D, Fig. S12A). While homozygous *tpte* mutants (*tpte*^*-/-*^) do not show evident developmental defects (Fig. S12B-E), they are unable to successfully regenerate their fins (Fig. 4I-J, Fig. S12E-G), similar to *lof*^-/-^ phenotypes (Fig. 3I-J). Strikingly, within the first 11h post-injury, we find only the first proliferative peak (1-3 hpa) to be absent (Fig. 4E-H, Fig. S12I-J). Furthermore, the Ca^2+^ injury responses remain unaffected in *tpte*^*-/-*^ (Fig. S12H). These observations suggest that loss of VSP-mediated electrical signal transduction in *tpte*^*-/-*^ is sufficient to induce regenerative defects at 5dpf. Importantly, tissue depolarisation by KCl treatment in uncut *tpte*^*-/-*^ does not result in increased proliferation (Fig. 4K vs. Fig. 3M, Fig. S12K). Hence, we conclude that VSP specifically senses and converts the V_mem_ depolarization signal into cell proliferation in the fin.

Finally, to understand whether modulating VSP sensitivity can trigger organ growth independently of injury, we generated conditional heat-shock transgenics that overexpress a modified version of VSP (VSP OE, Methods), resulting in increased phosphatase activity and quick response to V_mem_ fluctuations (*63*). Remarkably, overexpressing this form of VSP in uncut conditions (Fig. 4L, Fig. S13E-G, Methods) results in longer fins at 5 dpf (Fig. 4M, Fig. S13A,D), presenting a similar size increase to KCl-treated fins (Fig. 3K). Analogously, overexpressing VSP in injured fins also leads to successful regeneration at 3 dpa (Fig. S13A-C). Overall, these experiments show that VSP is the molecular effector that translates membrane depolarisation signals into cell proliferation, being necessary and sufficient to ensure complete tissue regeneration.

## Discussion

Our work reveals that the earliest organ-scale wound responses are closely instructed by the intrinsic bioelectrical state of tissues. Tissue-scale depolarization occurs immediately upon organ injury, induces proliferation and, consequently, fast-tracks organ regeneration. Such V_mem_ changes are sensed by a voltage-sensing protein, VSP, that converts electrical signals into intracellular chemical responses, resulting in controlled proliferation. This coupling primes the injured organ for successful regeneration.

By using sub-second-resolved live imaging, coupled with precise injuries in zebrafish larvae, we measured unprecedented tissue-scale V_mem_ (Fig. 1G-J) and Ca^2+^ (Fig. 1C-F) dynamics that are immediate tissue injury signals (100ms). Tissue electric dynamics often are modelled using equations borrowed from single neuron dynamics (*13, 65, 66*). Here, in contrast, by developing an electro-diffusive model that couples electric potential and ionic changes in the fin (Fig. 2A-D), we captured multi-timescale dynamics of early injury signals, inaccessible in previous frameworks. Our model uncovers two timescales occurring in this context: a millisecond response under quasi-static ionic concentration profiles, due to voltage short-circuiting caused by injury; and a second response, occurring in the seconds after, that is derived from the electrochemical interplay of ion diffusion and transport across the epithelium. Together, these two timescales quantitatively explain the ionic root of the post-injury electrical and intracellular Ca^2+^ signals, whose wound origin was thus far unresolved.

Interestingly, despite both fin epithelial layers becoming depolarized by the injury, we found that depolarization-induced proliferation occurs solely in the apical epithelial layer, due to VSP expression (Fig. 4A). Controlled proliferation upon depolarisation could indeed act as a general mechanism affecting organ size, as even in the absence of injury, longer fins were observed (Fig. 3L). This directly links tissue-level membrane potential changes to activation of proliferation, independently of Ca^2+^ signals. While the layer-specific expression of VSP remains enigmatic, understanding whether its location is restricted to tissues that are injury-prone will likely be relevant to enhance the proliferative capability of tissues with low regenerative potential.

Our data further shows that VSP directly transduces electrical signals into cell proliferation, as early as 1h post-injury (Fig. 4E-G). Previous *in vitro* studies have linked membrane depolarisation to proliferation through K-ras signalling (*58*). Future work addressing MAPK spatiotemporal dynamics subsequent to VSP activation will enable a closer investigation of the reported depolarization-induced signal integration in injured fins. While here we focused on the early proliferation dynamics controlled by electrical signals at 1h, we also observed that at 7h post-injury, cells proliferate independently of the described voltage-sensing mechanism. Given the timescale, this second proliferative peak is consistent with transcriptional activation of pro-regenerative signalling (*57, 67*).

Finally, our data suggests that enhancing K^+^ efflux compromises fin regeneration, by impairing proliferation triggered at 1 hour post-injury (Fig. 3G-H). A working hypothesis is that the applied manipulations (*lof*^*-/-*^ and FK506) increase K^+^ leakage, lowering the resting membrane potential of the tissue, hyperpolarizing V_mem_ (*20*). Given this lower V_mem_ level, upon wounding, the depolarization induced is not sufficient to trigger VSP conformation change, known to occur at a threshold of ∼40 mV (*59, 62, 63*). Interestingly, *longfin*^*t2*^ mutations and FK506-treatments have been found to promote fin outgrowth in adult zebrafish and other fishes (*20, 21, 25, 26*). This apparent discrepancy with our larval fin regeneration findings (Fig. 3G-J; S10) could be due to differences in the cell types that respond to injury: our larval system regenerates by essentially promoting epithelial proliferation through VSP activation (Fig. 4B,F,I-J), instead of a classic dedifferentiated, proliferative mesenchymal blastema structure (*20, 56, 68*). Deciphering the electrochemical dynamics of V_mem_ and ion fluxes in the adult mesenchymal blastema, especially considering the timescale of that regenerative process (2 weeks), is an exciting frontier to be explored.

Building on recent discoveries of bioelectrical processes at the heart of critical cellular and organismal functions (*66, 69*–*71*), our work suggests electrical regulation is a prevalent *in vivo* mechanism for epithelial tissues and organs in the need of fast communication. Our study opens an avenue for resolving the spatiotemporal interplay between fast electrical signals and slower biochemical signalling underlying organ regeneration.

## Acknowledgements

We thank Jan Brugués, Anne Grapin-Botton, Meritxell Huch, Jonathan Rodenfels and Michele Marass for critical manuscript feedback. We thank Luke D. Lavis for providing Janelia Fluor 552-Halotag dye, Palina Trus for initial efforts on performing Calcium live imaging in zebrafish, Tommaso Bianucci and Lucas Ribas for help with the proliferation analysis pipeline, Fenja Gawlas and Philipp Rustler for help with generating Ca^2+^ transgenic lines at the University of Freiburg. We are grateful to the following MPI-CBG facilities: Light Microscopy, Technology Development Studio; Electron Microscopy, Transgenic Core, and Zebrafish Unit. E.N., J.L., N.B., S.K., C.P., and R.M. acknowledge funding from the Max Planck Society and the Deutsche Forschungsgemeinschaft (DFG, German Research Foundation) under Germany’s Excellence Strategy - EXC-2068-390729961 - Cluster of Excellence Physics of Life of TU Dresden. J.L. has been supported by the ELBE Postdoctoral Fellowships Program and a MSCA Postdoctoral fellowship. A.B.A acknowledges funding by the Baden-Württemberg Stiftung (Elite programme for Postdocs), and DFG grant EXC294 (BIOSS Centre for Biological Signalling Studies), while supported by Wolfgang Driever.

## Author contributions

This work is a tightly integrated effort where experiment and theory were closely linked, resulting in key contributions by E.N., J.L., A.S. and C.D.: J.L. and E.N. performed the experiments, A.S. and C.D. developed the theory. Additional contributions: J.L. established sub-second live imaging setup. E.N., N.B., and S.K. established the *tpte*^*cbg22*^ line. N.B. and S.K. established the Tg(Hsp70:CiDr-VSP-L223f-mCherry)^*cbg23Tg*^ line. A.B.A. established the Tg(ubb:GCaMP6f)^*m1299*^ line. E.N. and S.K. performed cloning. C.P. helped obtain *lof*^*t2*^ datasets. F.J. and R.M. developed the project. E.N., J.L., A.S., C.D., F.J. and R.M. wrote the manuscript. All authors read and approved the final version of the manuscript.

## Data and materials availability

Source data for all main and supplementary figures are deposited on Edmond: doi.org/10.17617/3.6WBOSZ. Custom code for analysis pipelines and simulations is available on Zenodo (doi.org/<u>10.5281/zenodo.15019089, doi.org/10.5281/zenodo.15038032, doi.org/10.5281/zenodo.15037932</u>). Plasmids and zebrafish lines generated in this study are available upon request.

