## Supplemental_Materials for "Injury-induced electrochemical coupling triggers regenerative cell proliferation"

**The PDF file includes:**

Figs. S1 to S18
Legends for Movies S1 to S9
Materials and Methods
Supplementary Theory Notes
References

**Other Supplementary Materials for this manuscript include the following:**

Movies S1 to S9

**CONTENTS**


Supplementary figures 4

Supplementary movie legends 19

Materials and Methods 21

Experimental procedures 21

1. Ethics Statement 21
2. Zebrafish lines and maintenance 21
3. $tpte^{cbg22}$ mutant 21
4. Transgenic lines generation 22
5. Genotyping 23
6. Voltron2 Cloning, mRNA synthesis and embryo microinjection 24
7. GAP27 peptide heart injections 24
8. Chemical treatments 25
    1. Voltron2 labelling with JF552 25
    2. $Ca^{2+}$ free E3 medium 25
    3. KCl supplement 25
    4. FK506 26
9. Spinning disk live imaging and UV-Laser microdissection 26
10. Embryo heat-shocks 27
11. Mechanical amputation and injury 27
    1. Fin fold regeneration assay 28
    2. Stereoscope $Ca^{2+}$ imaging following fin injury 28
12. Proliferation assay 28
13. Immunofluorescence 28
14. Transmission electron microscopy (TEM) 29

Quantitative procedures 30

1. Epithelial cell segmentation and registration 30
2. GCaMP6f cytosolic signal quantification 31
3. Voltron2:JF552 membrane signal quantification 33

4. Fin and body length quantifications 34
5. Fin epithelial layer thickness and interstitial space quantification 34
6. Proliferation quantification 35
7. VSP intensity quantification 36

Supplementary Theory Notes 36

I. Multi-compartment model of ion transport 36
1. Model and geometry 36
2. Ion transport and diffusion 37

II. Multi-timescale electrical response to injury 38
1. Electro-diffusive dynamics in the interstitial fluid 38
2. Electro-diffusive dynamics in the epithelium 40

III. Comparison of the theory with the experiment 40
1. Fast changes in electric potential 40
2. Cellular calcium response 41
3. Calcium activation wave in the epithelium 42
4. Cell-cell variability of the calcium activation threshold can explain noisy calcium activation wave 43

IV. Effect of adding potassium chloride to the external medium on the ion electro-diffusive dynamics 45
1. Effect of potassium chloride supplement on epithelial electric potential 45
2. Slower diffusion of sodium ion in the presence of potassium ion 46

V. Diffusion front and effective diffusion constant 48
1. Simple diffusion with threshold 48
2. Effective diffusion constant in the tissue model 48

VI. Parameters used in the figures and movie 49

References 50

### SUPPLEMENTARY FIGURES

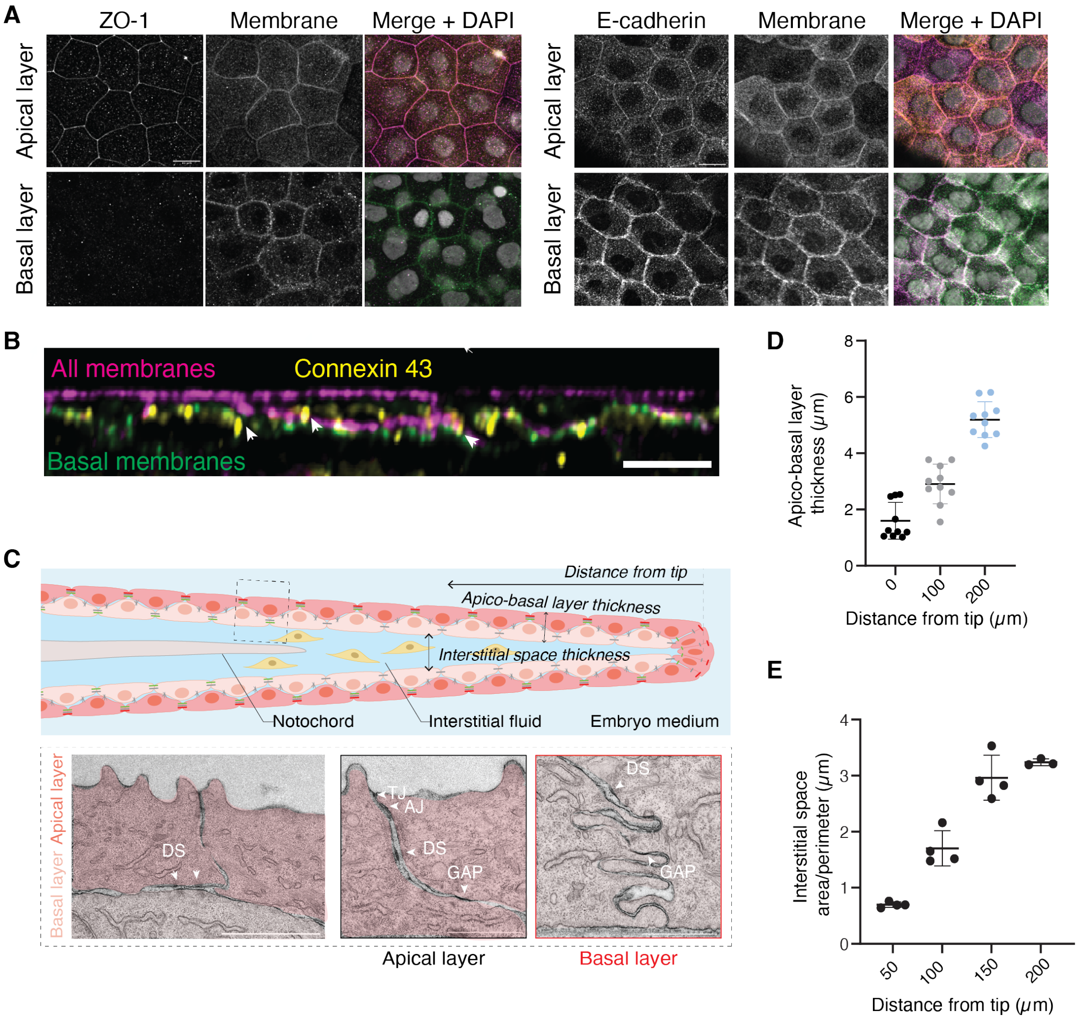


FIG. S1: **Zebrafish caudal fin at 2 days-post-fertilization displays planar apico-basal epithelial tissue architecture.** (A) Confocal images of immunostainings against tight junction protein (ZO-1, left) and adherens junction protein (E-cadherin, right) in apical and basal fin epithelial layers. (B) XZ view of lattice light sheet image of 2dpf fin immunostained against gap junction proteins (Connexin 43, yellow, white arrowhead). All membranes of epithelial cells, magenta; membranes of basal epithelial cells, green. (C) Top, schematic of fin architecture, with dimensions measured in D and E. Dashed box indicates position of the fin epithelium at the notochord level. Bottom, electron transmission micrographs of 2dpf fin apical and basal epithelial layers at notochord level (160 $\mu$m from fin tip). White arrows: tight junctions (TJ), adherens junctions (AJ), desmosomes (DS) and gap junctions (GAP). Basal epithelial layer only contains gap junctions at the notochord, $\sim$160 $\mu$m from the fin tip. (D) Thickness of double epithelial layer (apical and basal epithelial cells) along fin positions from its distal tip. (E) Fin interstitial space area-to-perimeter ratio, quantified from XZ images in (B) vs. distance from fin tip. Typical injury site is at 50$\mu$m from the fin tip. Scale bar: 10 $\mu$m (A), 20 $\mu$m (B), 2 $\mu$m (left), 1 $\mu$m (right) (C).

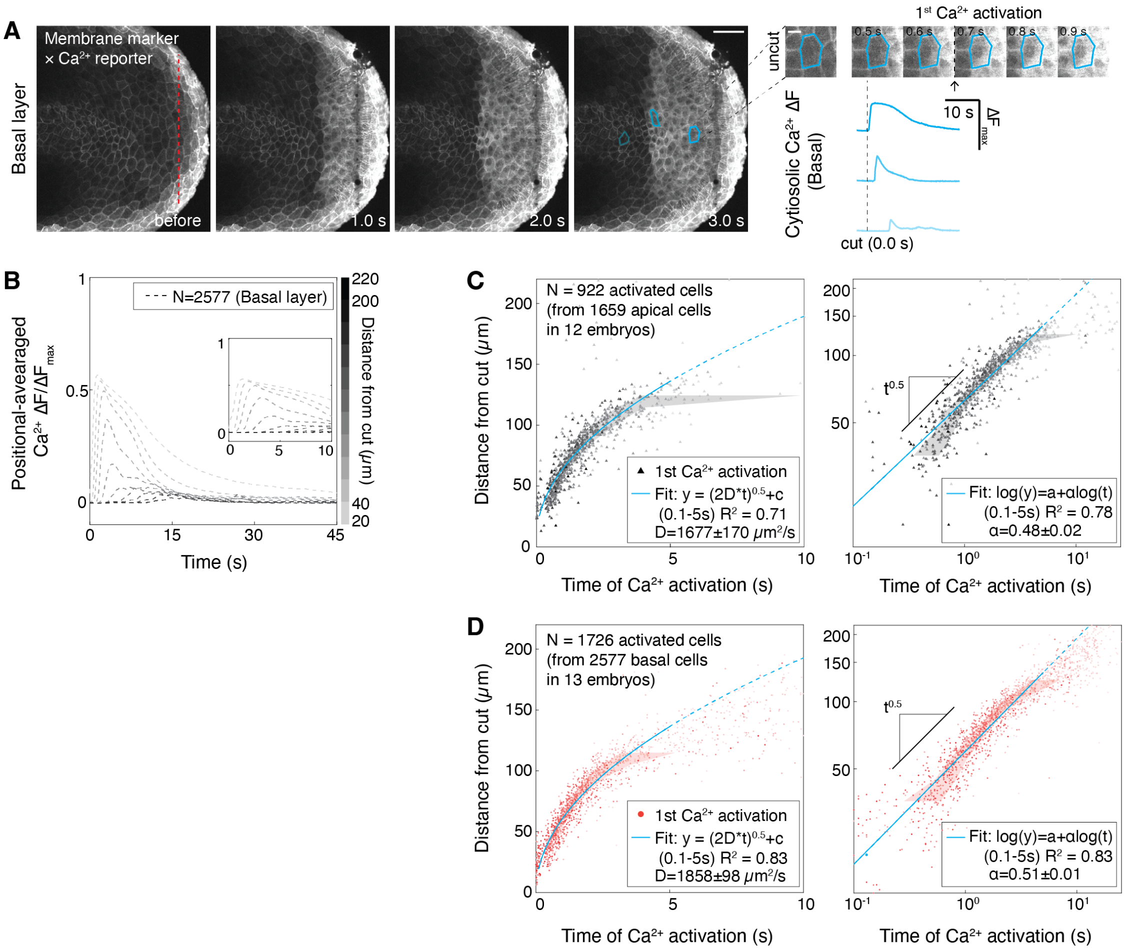


FIG. S2: **Fast intracellular Calcium wavefront propagates and decays concomitantly in apical and basal epithelial layers.** (A) 2dpf fin before and after cut (red dash), in transgenics labeling intracellular Calcium (ubb:GCaMP6f) and basal epithelial layer membranes (tp63:CAAX-GFP) (see Movie S1). Right, time-lapse images of a representative individual cell and changes of cytosolic $Ca^{2+}$ intensity (ΔF, vs uncut). (B) Position-averaged changes of cytosolic $Ca^{2+}$ intensity ($\Delta F/\Delta F_{max}$, vs uncut, normalized per embryo) from basal fin epithelium vs. time. Inset: zoom-in view during 0-10 s. 13 larvae analysed. (C-D) Position of apical (C, from Fig.1F) or basal (D) epithelial cells relative to wound vs. time of $Ca^{2+}$ wavefront activation. Right, log-to-log scale. Shade, 25%-75% percentile of single cell data. Cyan lines, diffusive fit in linear scale (D: diffusion coefficient), linear fit ($\alpha$: scaling exponent) in log-to-log scale (0.1s to 5s). Cyan dashed, continuation of diffusive fit in linear scale (0.1s to 20s) or log-to-log scale. Error, 95% CI of parameter fits. R2, goodness of fit. N: number of cells (B,C,D).

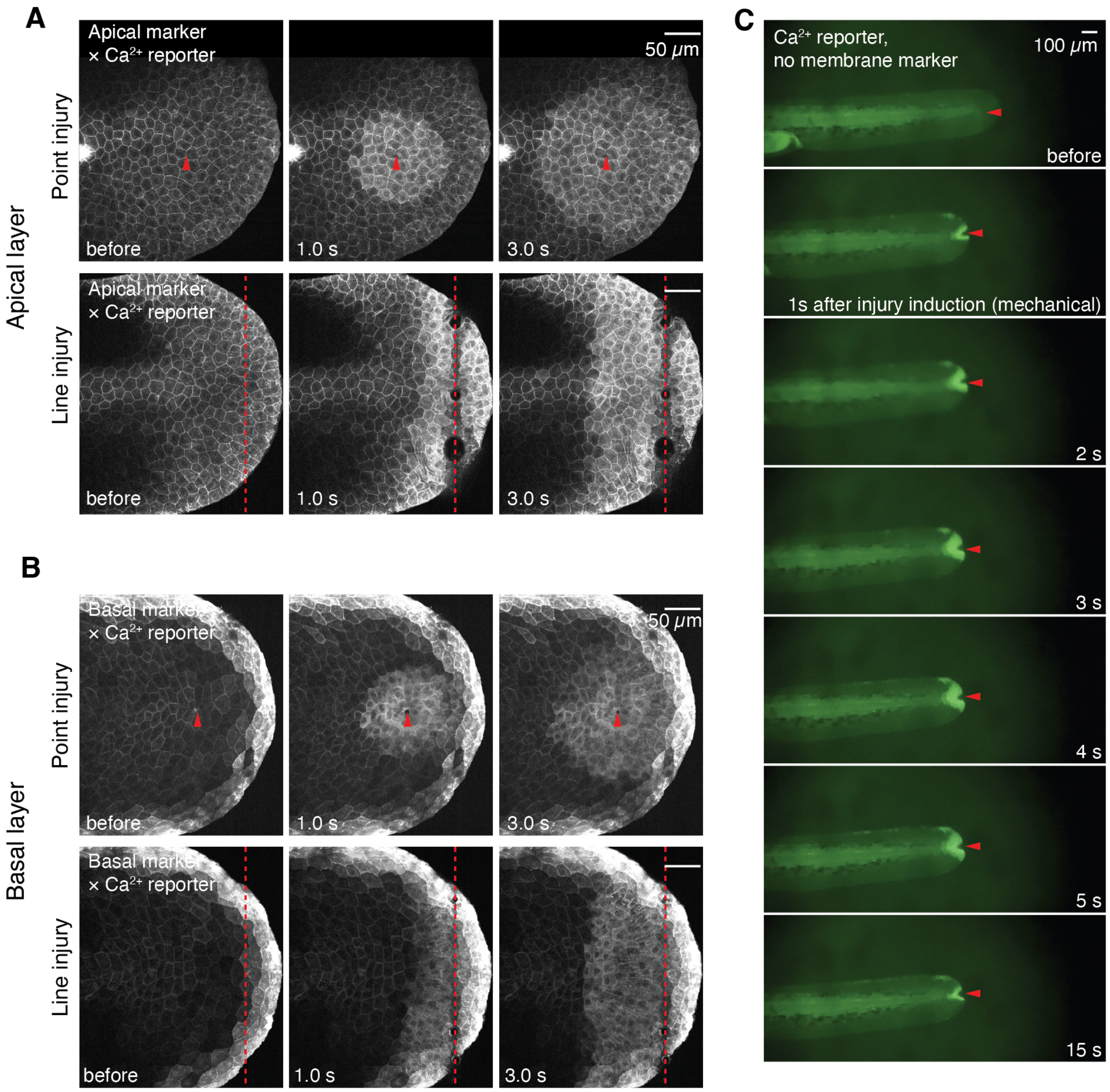


FIG. S3: **Formation of the Calcium wavefront is independent of injury size or injury method.** (A) Time-lapse images of 2dpf fin before and after a point-shaped (red arrowhead) or line-shaped UV-laser cut (red dash) in double transgenics labeling intracellular $Ca^{2+}$ (ubb:GCaMP6f) and apical epithelial layer membranes (top, claudinb:Lyn-GFP)(see Movie S2). Scale bars: $50\mu m$. (B) Time-lapse images of 2dpf fin before and after a point-shaped (red arrowhead) or line shaped UV-laser cut (red dash), in double transgenics labeling intracellular Calcium (ubb:GCaMP6f) and basal epithelial layer membranes (bottom, tp63:CAAX-GFP) (see Movie S2). Scale bars: $50\mu m$. Note that in (A) and (B), time-lapse imaging before and after UV-laser ablation were executed as separate experimental blocks (Methods). (C) Time-lapse images of 2dpf fin before and after mechanical cut (red arrowhead), in transgenic line labeling intracellular Calcium (ubb:GCaMP6f) (see Movie S3). Scale bar: $100\mu m$.

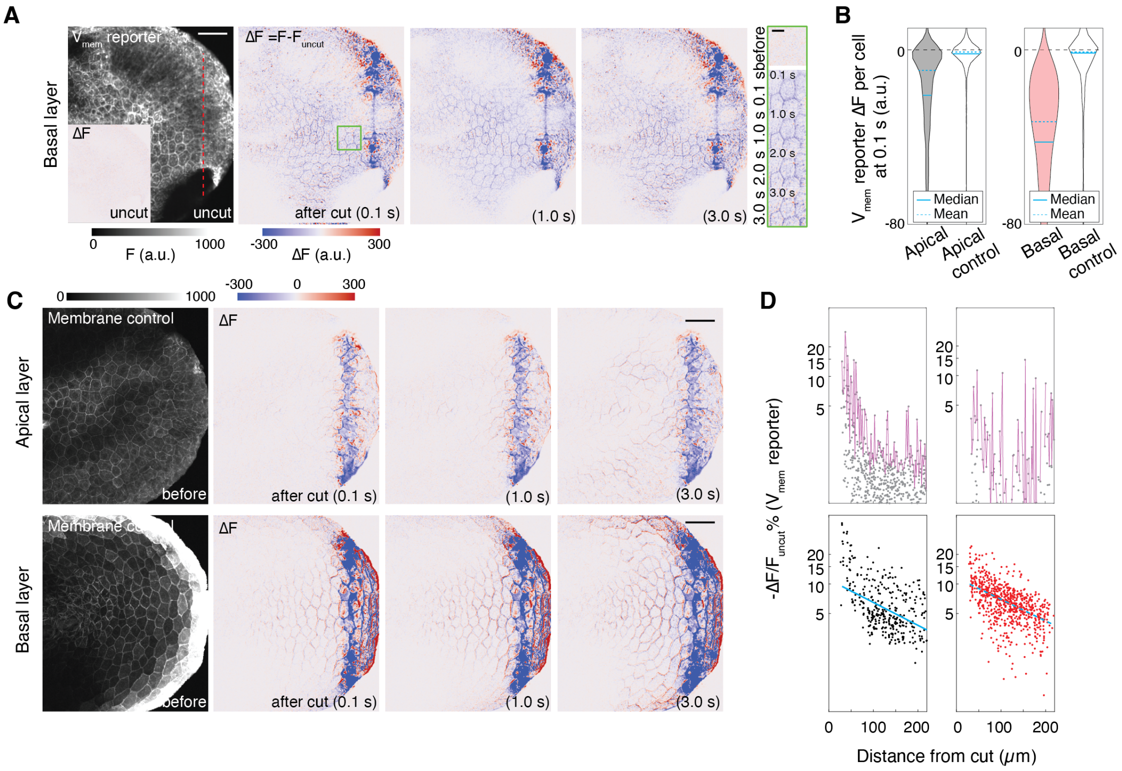


FIG. S4: **$V_{mem}$ reporter dynamics upon fin injury.** (A) Left, 2dpf basal fin epithelium (tp63:CAAX-GFP) labelled with Voltron2 ($V_{mem}$ reporter) before cut (red dash). Inset, change of membrane Voltron2 intensity in uncut fins (ΔF, vs previous 0.1s uncut). Middle to right, change of membrane Voltron2 intensity at 0.1s, 1.0s and 3.0s post-cut (ΔF, vs uncut). Green inset, cellular detail. Scale bar: Left, $50\mu m$; Inset, $20\mu m$. (B) Change of intensity (ΔF, vs uncut) at 0.1s post-cut, compared between apical epithelial cells expressing Voltron2 (grey, N=9 larvae; 946 cells) or respective membrane marker control (Claudinb:Lyn-GFP, white, N=10 larvae; 1373 cells); and basal epithelial cells expressing Voltron2 (pink, N=9 larvae; 914 cells) or respective membrane marker control (tp63:CAAX-GFP, white, N=5 larvae; 180 cells). (C) Left, 2dpf control fin epithelium expressing apical (top, claudinb:Lyn-GFP) or basal (bottom, tp63:CAAX-GFP) membrane marker. Middle to right, change of membrane marker intensity at 0.1s, 1.0s and 3.0s post-cut (ΔF, vs uncut). Scale bars: $50\mu m$. (D) Top, noise floors (magenta) in the fractional drops of Voltron2 intensity (-ΔF/F0) vs. position relative to wound, constructed from the change of intensity of membrane marker controls at 0.1s after cut. Bottom, fractional drops of Voltron2 intensity (-ΔF/F0) vs. position relative to wound, for Voltron2-labelled apical and basal epithelial cells, post-noise floor extraction, at 0.1s post-cut.

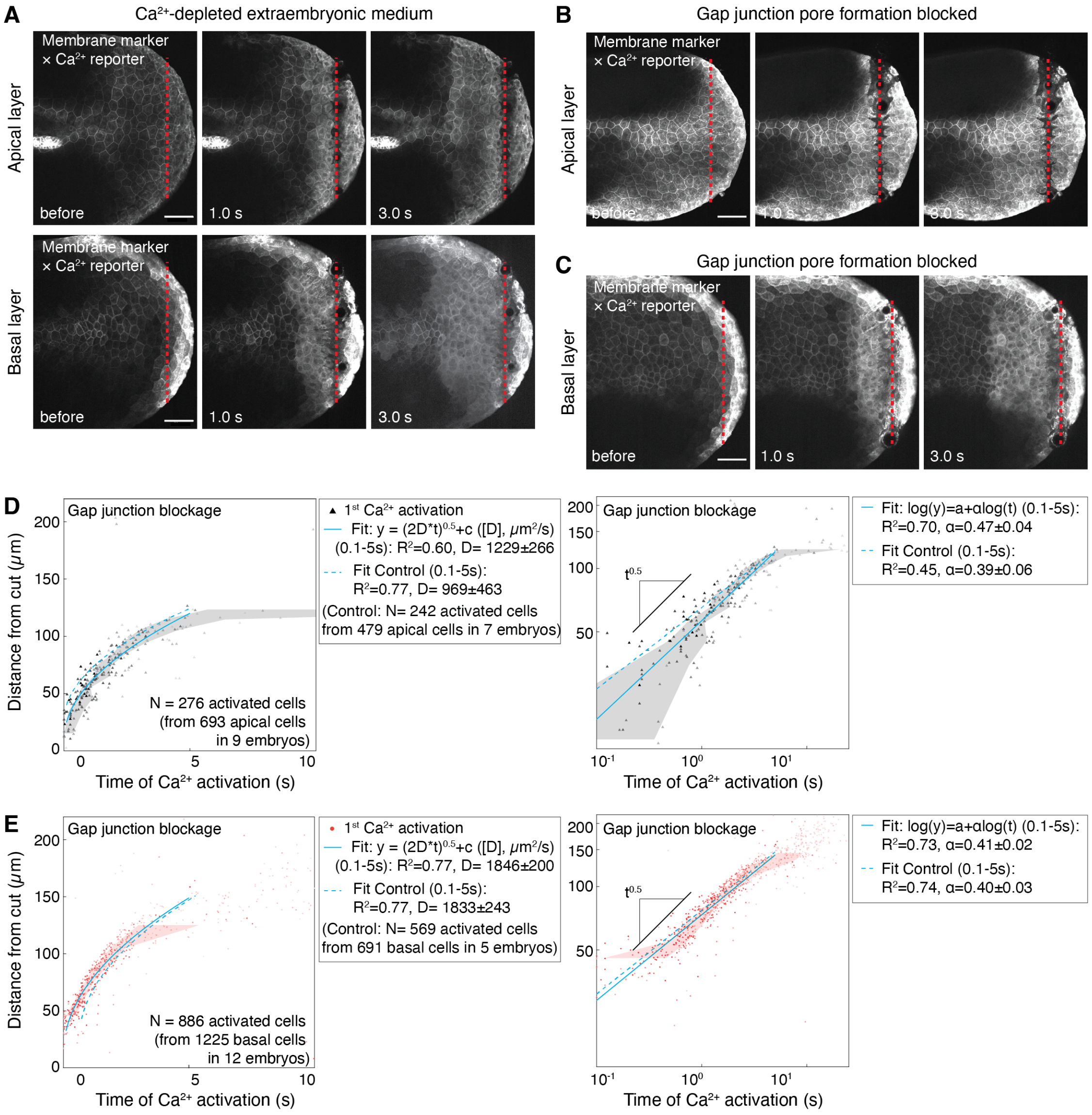


FIG. S5: **$Ca^{2+}$ wavefront propagation is not affected by extraembryonic $Ca^{2+}$ depletion nor gap junction blockage.** (A) Calcium wavefront dynamics in larvae incubated $Ca^{2+}$ free medium (Methods). Top, Time-lapse images of apical fin epithelium labelled with intracellular $Ca^{2+}$ reporter (double transgenics, claudinb:Lyn-GFP and ubb:GCaMP6f). Bottom, Time-lapse images of basal fin epithelium labelled with intracellular $Ca^{2+}$ reporter (double transgenics, tp63:CAAX-GFP and ubb:GCaMP6f). See Movie S5. (B) Calcium wavefront dynamics in apical fin epithelium with GAP junction pore formation inhibited (double transgenics, claudinb:Lyn-GFP and ubb:GCaMP6f). Note that time-lapse imaging before and after UV-laser ablation was executed as separate experimental blocks (Methods). See Movie S6. (C) Calcium wavefront dynamics in basal fin epithelium with GAP junction pore formation inhibited (double transgenics, tp63:CAAX-GFP and ubb:GCaMP6f). See Movie S7. *(Continued on next page)*

(D-E) Position of apical (D) and basal (E) epithelial cells relative to wound vs. time of $Ca^{2+}$ wavefront (triangle) activation in gap junction-blocked condition. Right, log-to-log scale. Shade, 25%-75% percentiles. Cyan lines, diffusive fit in linear scale and linear fit in log-to-log scale (0.1s-5s). Cyan dash, fit (0.1s-5s) from respective control datasets (phenol red injections). All scale bars: $50\mu m$.

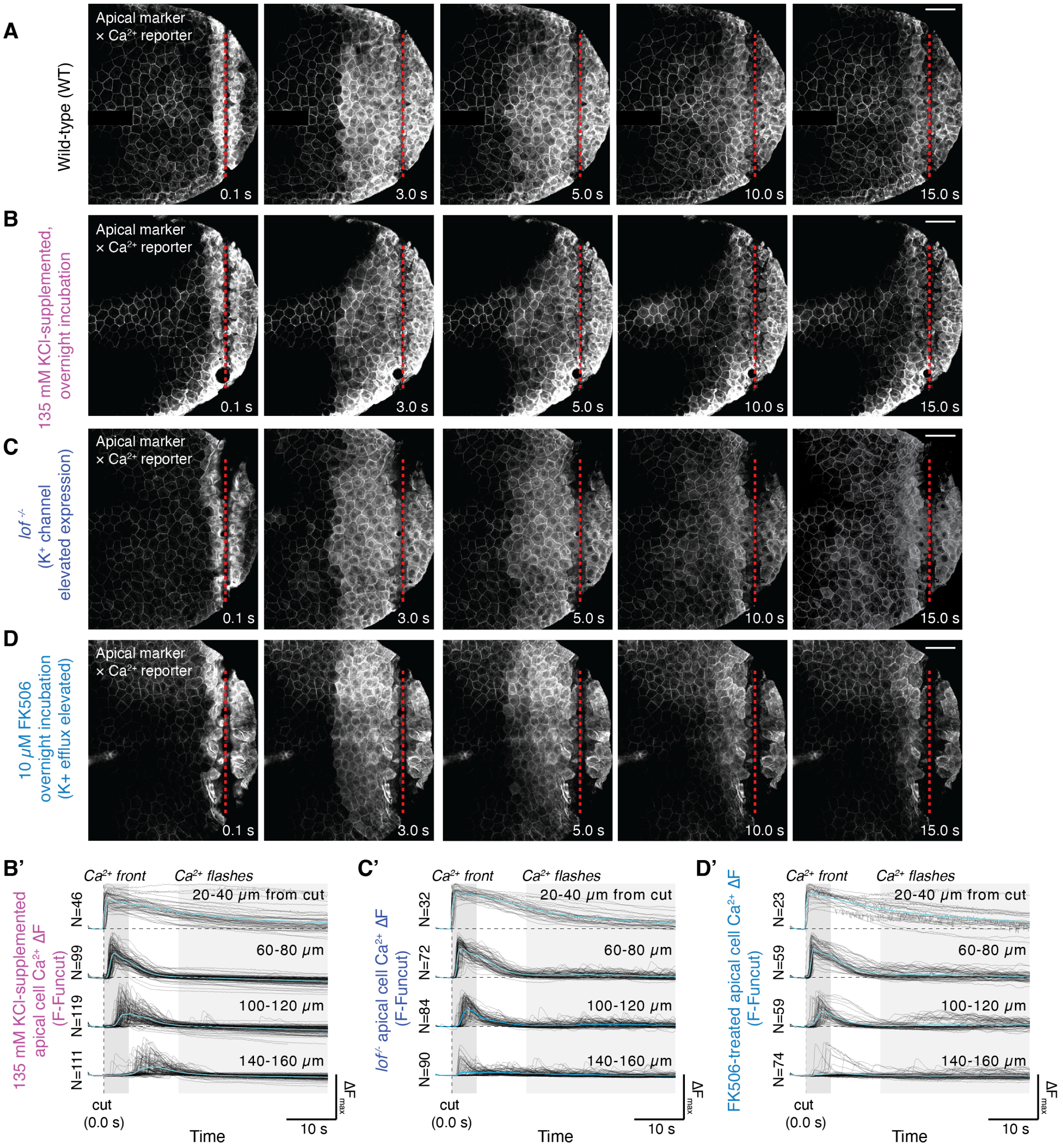


FIG. S6: **Impact of K$^+$ flux manipulations on fin Ca$^{2+}$ dynamics upon injury.** (A-D) 2dpf fins at 0.1s, 3s, 5s, 10s and 15s after fin cut (red dash), in double transgenics labeling intracellular Calcium (ubb:GCaMP6f) and apical epithelial layer membranes (claudinb:Lyn-GFP). (A) wildtype; (B) KCl-treated larvae; (C) *lof*$^{-/-}$; and (D) FK506-treated larvae. (B') Change of KCl-supplemented cytosolic Ca2+ intensity (ΔF, vs uncut) for individual cells (gray) and positional averages (cyan) vs. time. (C') Change of *lof*$^{-/-}$ cytosolic Ca2+ intensity (ΔF, vs uncut) for individual cells (gray) and positional averages (cyan) vs. time. (D') Change of FK506-treated cytosolic Ca$^{2+}$ intensity (ΔF, vs uncut) for individual cells (gray) and positional averages (cyan) vs. time. Scale bars for all images: 50$\mu m$. (See Movies S8-S9)

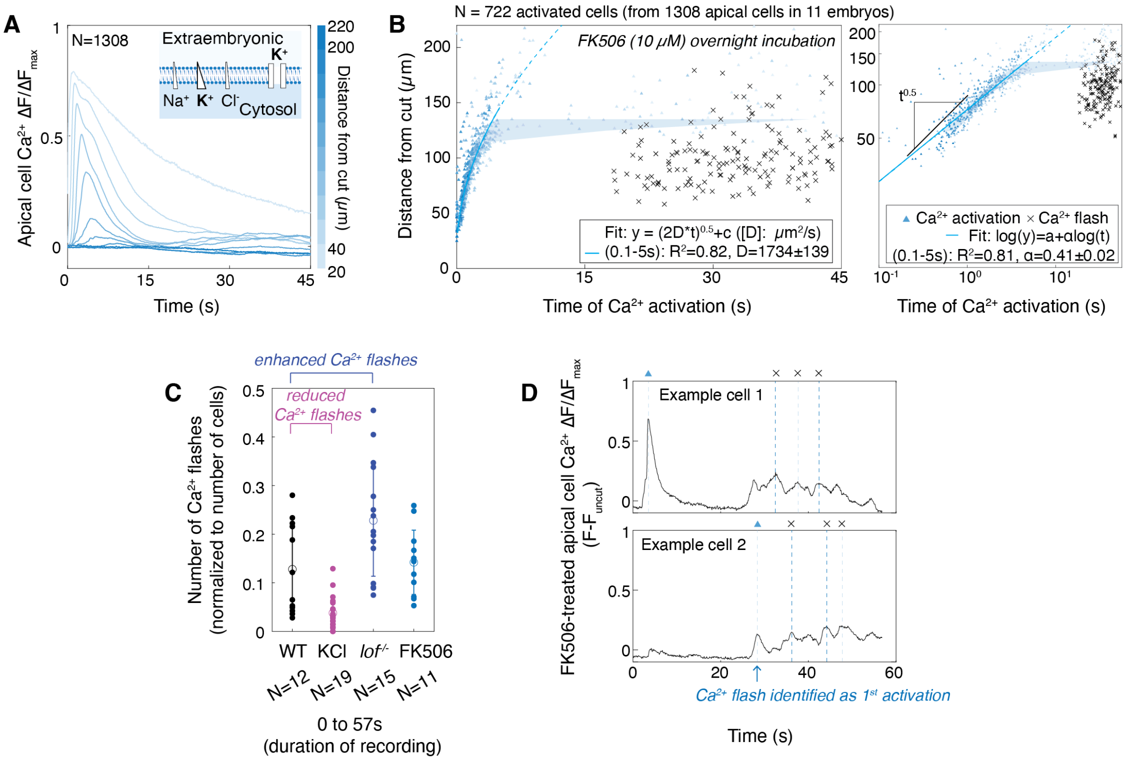


FIG. S7: **Quantification of injury $Ca^{2+}$ dynamics under $K^+$ flux manipulations.** (A) Position-averaged changes of normalized cytosolic $Ca^{2+}$ intensity ($\Delta F/\Delta F_{max}$, vs uncut, normalized per embryo) from apical fin epithelium vs. time, in FK506-treated larvae. Inset: Depiction of ionic concentration (blue) and transport across compartments in FK506-treatments. (B) Position of apical epithelial cells relative to wound vs. time of $Ca^{2+}$ wavefront (triangle) or flashes (cross) activation, in FK506-treatments. Shades, 25%-75% percentile. Cyan lines: diffusive fit in linear scale; linear fit in log-to-log scale (0.1s - 5s). Cyan dash, continuation of diffusive fit in linear scale (0.1s - 20s). Error, 95% CI of parameter fits. R2, goodness of fit. (C) Number of $Ca^{2+}$ flash events resulting from fin injury with UV-laser ablation (normalized by number of cells per embryo), across the K+ manipulation conditions used in this study. Mean +/- SD. N, number of larvae. (D) Two examples of fin apical cells under FK506-treatment, with identified $1^{st}$ $Ca^{2+}$ activation (triangle). Note that $1^{st}$ $Ca^{2+}$ activation (triangle) usually corresponds to a wavefront (example 1), with $Ca^{2+}$ flashes (cross) activated subsequently. In FK506-treated larvae (Fig. S6C-D) as well as *lof*$^{-/-}$ (Fig. 2J), the $1^{st}$ $Ca^{2+}$ activation (triangle) sometimes corresponds to a flash event (cross) instead of a wavefront (example 2). This accounts for the $Ca^{2+}$ flashes under FK506 treatment are being as prominent in Fig. S7C, but can be clearly observed in Fig. S7B (compared to wt, Fig. 1F) and Movie S9.

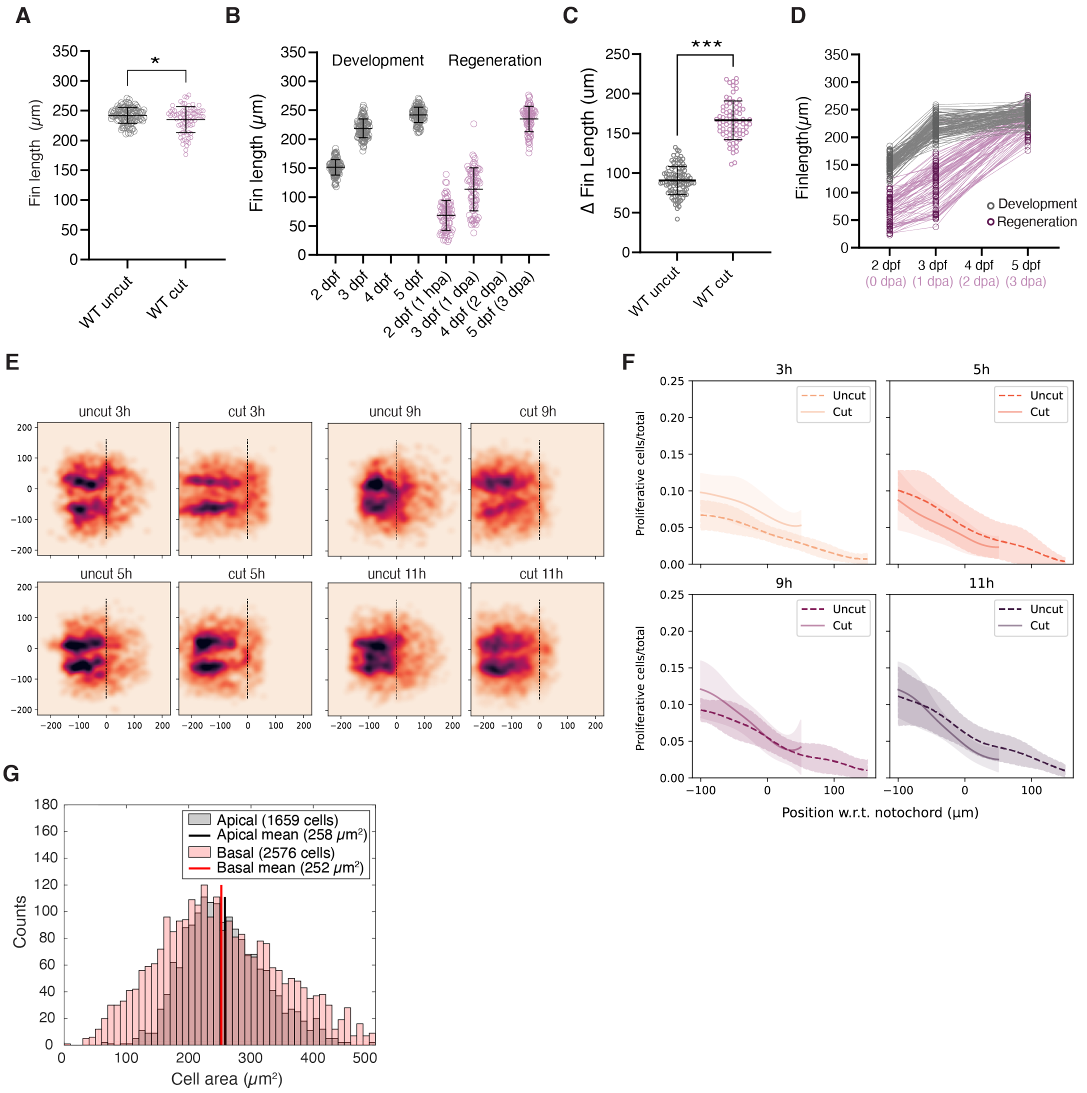


FIG. S8: **Fin length and proliferation dynamics in development and regeneration.** (A) Fin length measurements in 5dpf WT uncut vs regenerated at 3dpa. (B) Fin lengths in development (grey) and regeneration (pink), between 2-5 dpf. (C) Fin growth difference in length between 2-5 dpf in WT uncut vs cut (from B). (D) Time course of fin length from individual embryos in development (grey) and regeneration (pink) (from B). E) Heatmaps displaying probability of occurrence of a proliferative cell within fin area at 3h, 5h, 9h and 11h. Dashed line, notochord tip. (F) Fraction of proliferative cells vs. position at 3h, 5h, 9h and 11h uncut (dash line) and cut (full line). Zero indicates notochord. Shaded areas, SD. (G) Distribution of cell area measurements in apical and basal epithelial cells, in 2 dpf fins. A,B,C: Mean +/- SD. A-D: n = 101 embryos for development, 76 embryos for regeneration. E-F: n = 15 embryos per condition. All statistics: *p≤0.05, ***p≤0.001; two-tailed, unpaired, non-parametric Mann-Whitney tests.

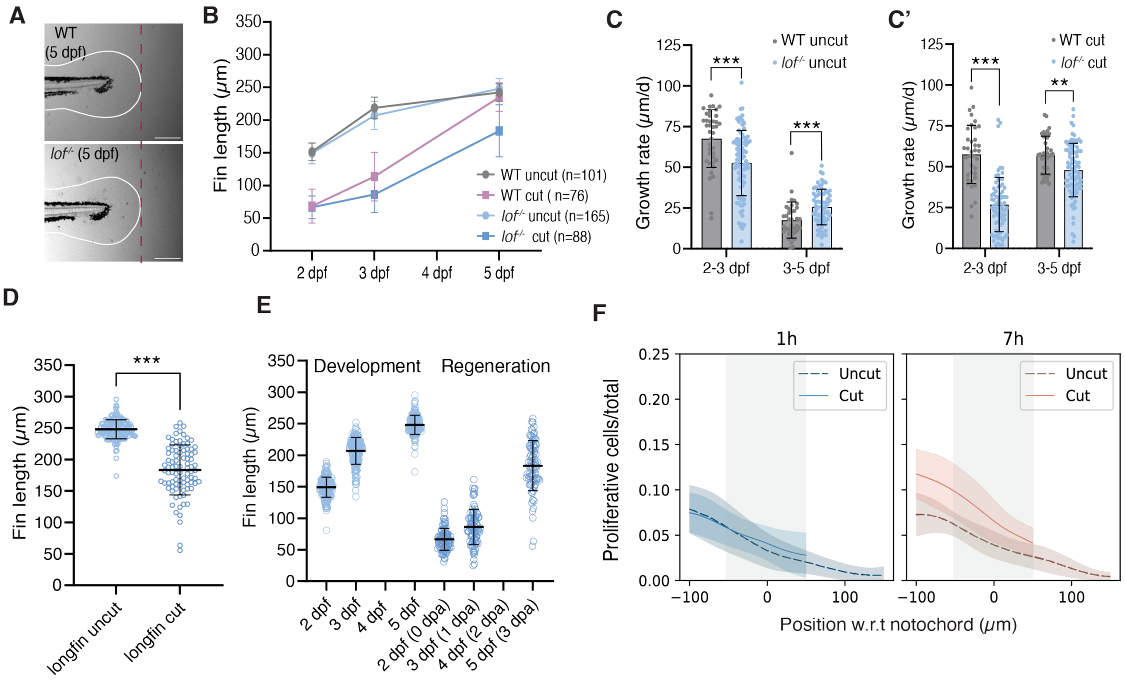


FIG. S9: **Increased $K^+$ leakage affects proliferation at 1hpa and impairs fin length regeneration**. (A) 5dpf WT and *lof*$^{-/-}$ uncut fins. Scale bar 200$\mu$m. (B) Fin lengths in uncut and cut conditions in WT vs *lof*$^{-/-}$, from 2-5 dpf. Mean +/- SD. (C) Fin growth rates comparison between WT and *lof*$^{-/-}$ uncut and (C') cut, calculated as the difference in fin length between indicated timepoints, divided by time (data from B). (D) Fin length measurements in *lof*$^{-/-}$ uncut vs cut at 5dpf (from B). (E) Fin lengths in development and regeneration in *lof*$^{-/-}$, between 2-5 dpf (individual data from B). Mean +/- SD. (F) Fraction of proliferative cells vs. position at 1h and 7h uncut vs cut in *lof*$^{-/-}$ (from Fig. 3G). Zero, notochord tip. Grey area indicates 100$\mu$m from the wound. Shaded areas indicate SD. n, number of embryos. All statistics: **p≤0.01, ***p≤0.001, two-tailed, unpaired, non-parametric Mann-Whitney tests.

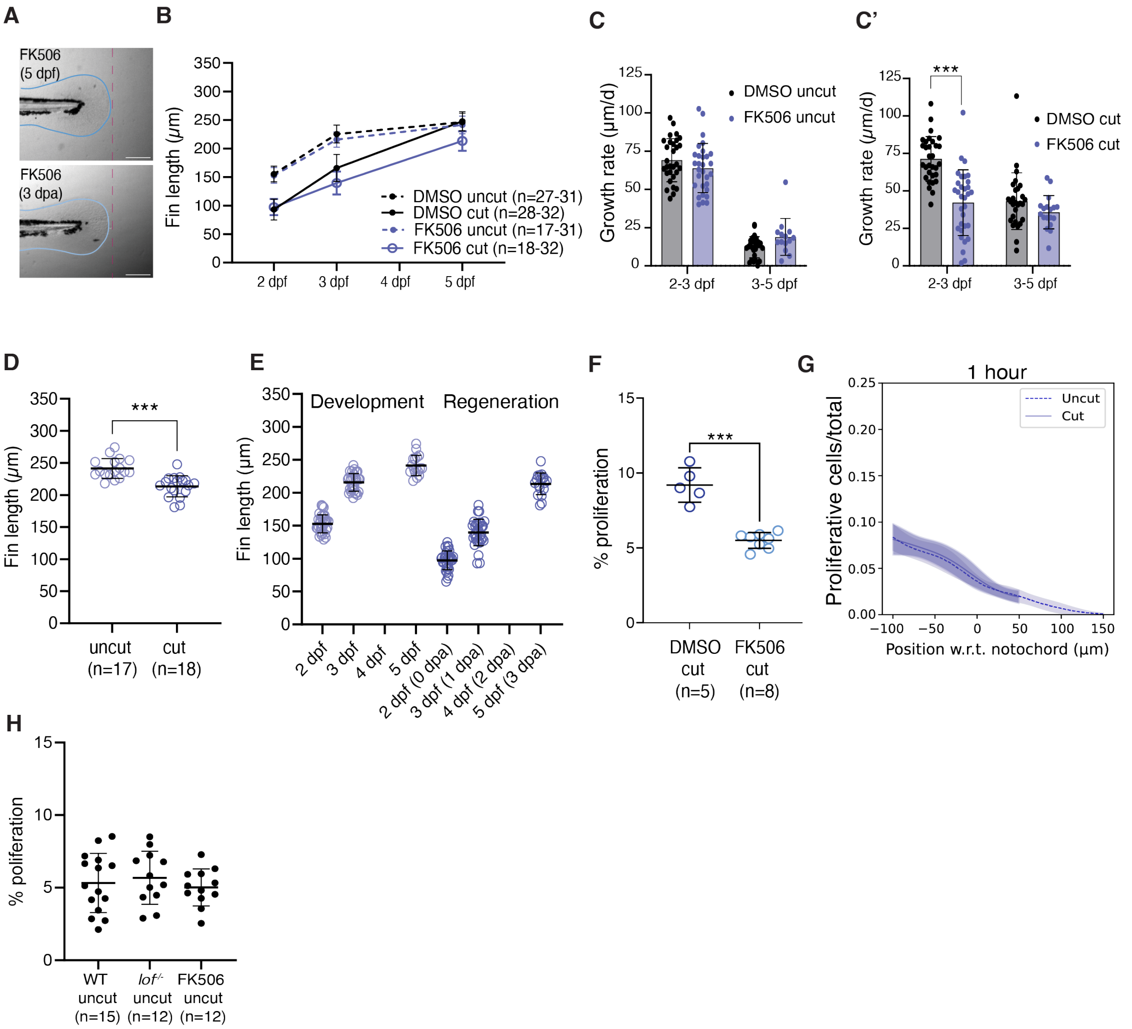


FIG. S10: **FK506 treatment mimics lof-/- mutants in proliferation and regeneration dynamics.** (A) WT and FK0506 treated uncut fins at 5 dpf. Scale bar, 200$\mu$m. (B) Fin lengths in uncut and cut conditions in WT vs FK506, from 2-5 dpf. (C) Fin growth rates comparison between WT and FK506 uncut and (C') cut, calculated as the difference in fin length between indicated timepoints, divided by time (from B). (D) Fin length measurements in FK506 uncut and cut at 5dpf (from B). (E) Fin lengths in development and regeneration in FK506-treated larvae, between 2-5 dpf. (F) Percentage of proliferative (EdU+) per total cells (DAPI+), in 1hpa cut fins, in WT vs FK506-treated larvae. (G) Fraction of proliferative cells vs. fin position inuncut vs cut fins (1h), in FK506-treated larvae. Zero, notochord tip. Line indicates average, shaded areas indicate SD. (H) Percentage of proliferative per total cells in 2dpf uncut fins (1h EdU) in WT vs *lof*$^{-/-}$ vs FK506. For B-F,H: mean +/- SD. All statistics: ***p$\leq$0.001, two-tailed, unpaired, non-parametric Mann-Whitney tests. n, number of embryos.

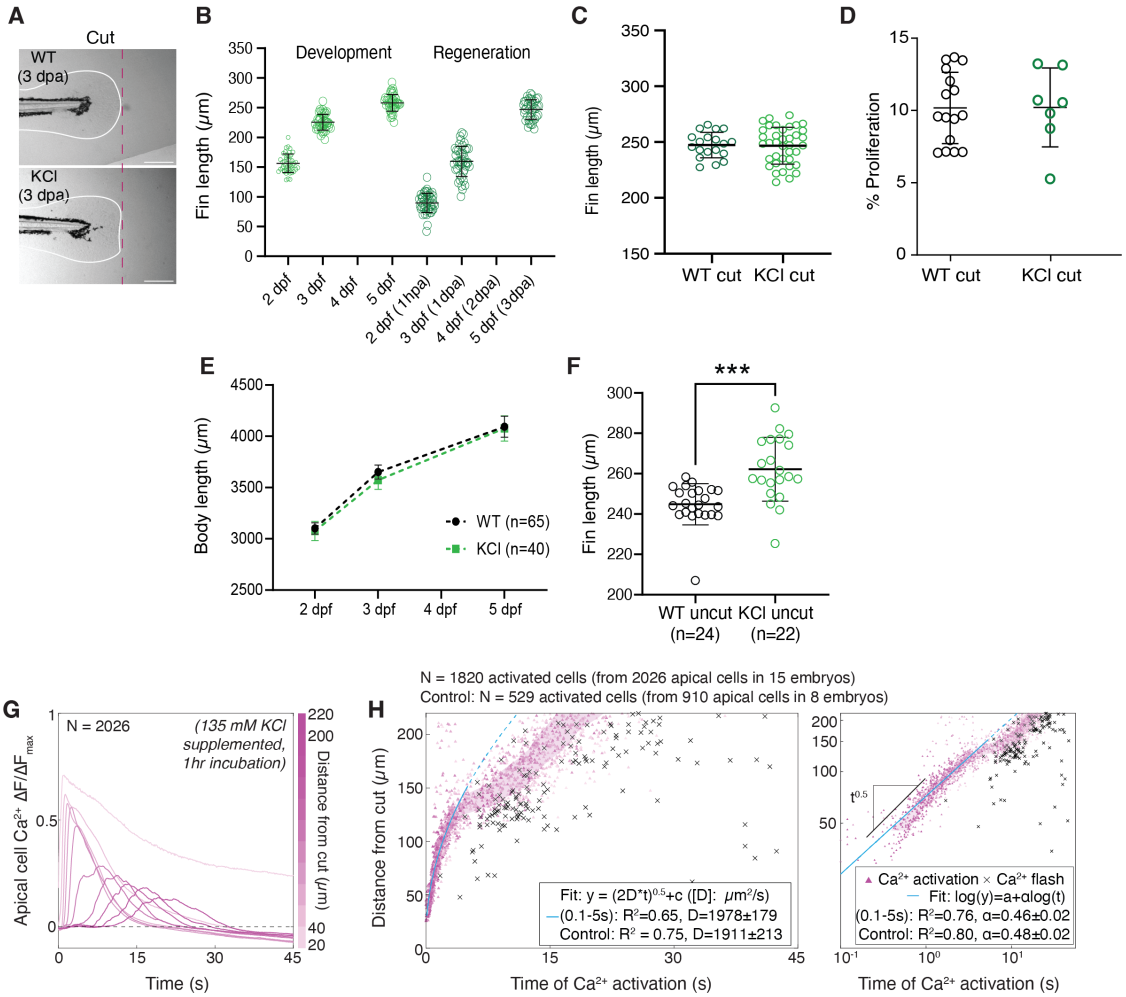


FIG. S11: **Fin regeneration and proliferation resemble wildtype dynamics upon KCl treatment.** (A) WT and KCl-treated regenerated fins at 5 dpf. Scale bar 200$\mu$m. (B) Fin lengths in development (light green) and regeneration (darker green) in 1h KCl treatment conditions, between 2-5 dpf. (C) Fin length measurements at 5 dpf regenerated fins (cut), in E3 vs KCl-treated (from B). (D) Percentage of proliferative per total cells in WT vs KCl conditions, at 1h post-cut. (E) Body length measurements in WT vs KCl-treated larvae between 2-5 dpf. (F) Fin length measurements at 5 dpf in E3 vs 12h KCl incubation. (G) Position-averaged changes of normalized cytosolic $Ca^{2+}$ intensity ($\Delta F/\Delta F$max, vs uncut, normalized per embryo) from apical epithelium vs. time, in KCl-treated embryos. (H) Position of apical epithelial cells relative to wound vs. time of $Ca^{2+}$ wavefront (triangle) or flash (cross) activation, in 1 hour KCl-treated larvae vs control (Methods). Right, log-to-log scale. Cyan lines: diffusive fit in linear scale; linear fit in log-to-log scale (0.1s to 5s). Cyan dash, continuation of diffusive fit in linear scale or log-to-log scale. n, number of larvae (B-F) or cells (G-H). All statistics: ***p≤0.001, two-tailed, unparied, parametric t-test.

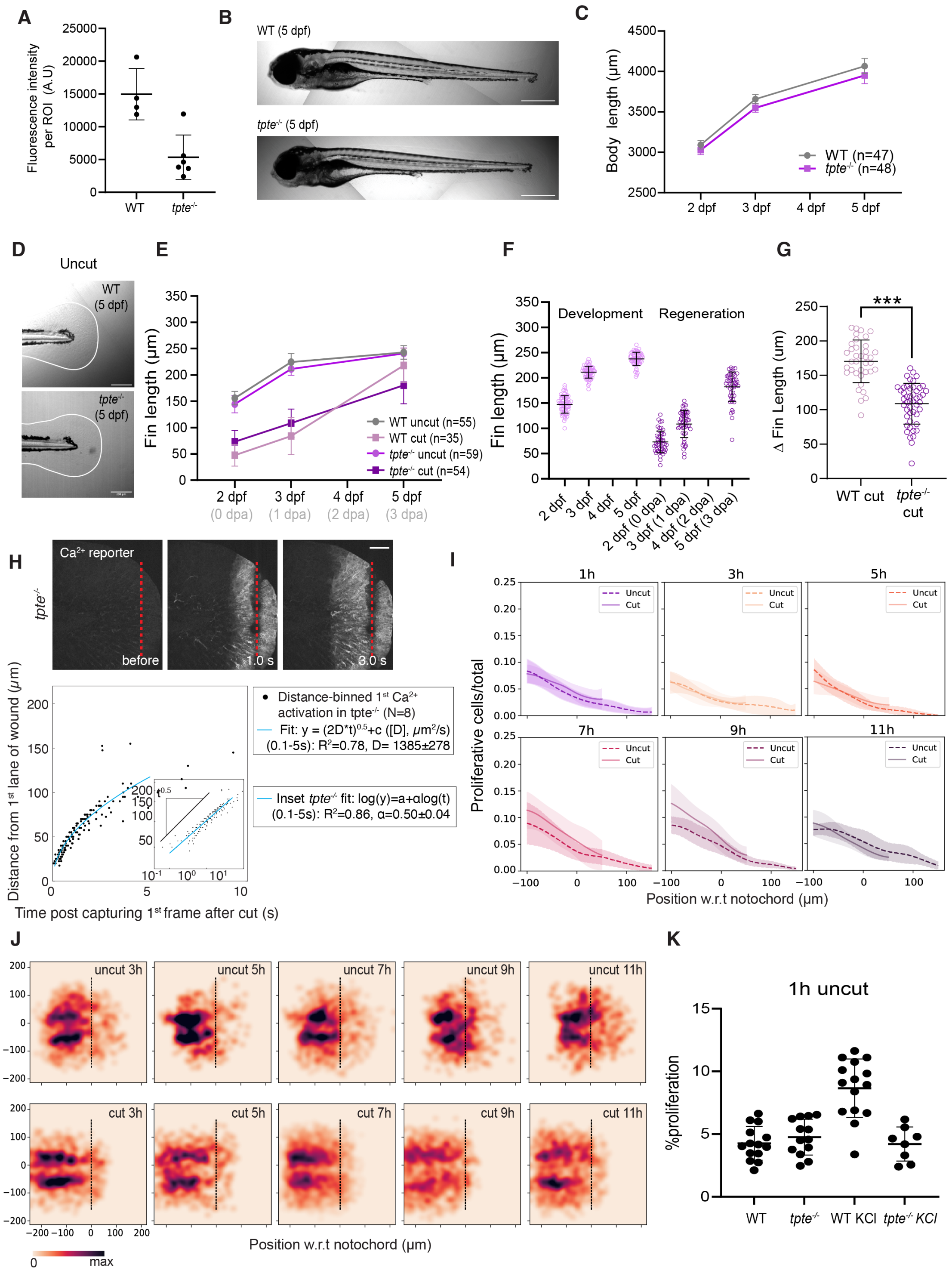
A
B
C
D
E
F
G
H
I
J
K
WT (5 dpf)
tpte-/- (5 dpf)
Uncut
Ca2+ reporter
before
1.0 s
3.0 s
Development
Regeneration
WT (n=47)
tpte-/- (n=48)
WT uncut (n=55)
WT cut (n=35)
tpte-/- uncut (n=59)
tpte-/- cut (n=54)
Distance-binned 1st Ca2+ activation in tpte-/- (N=8)
Fit: y = (2D*t)0.5+c ([D], μm2/s) (0.1-5s): R2=0.78, D= 1385±278
Inset tpte-/- fit: log(y)=a+αlog(t) (0.1-5s): R2=0.86, α=0.50±0.04
1h uncut
Fluorescence intensity per ROI (A.U)
Body length (μm)
Fin length (μm)
Δ Fin Length (μm)
Proliferative cells/total
%proliferation
Distance from 1st lane of wound (μm)
Time post capturing 1st frame after cut (s)
Position w.r.t notochord (μm)
WT cut
tpte-/- cut
WT KCl
tpte-/- KCl
0
max

FIG. S12: ***tpte***$^{-/-}$ **display impaired fin regeneration and associated proliferation, despite normal development.** (A) Intensity quantifications of VSP immunostainings, from mean intensity projections of both apical and basal epithelial layers in WT (n = 4 embryos) vs *tpte*$^{-/-}$ (n = 6 embryos). (B) Brightfield images of 5 dpf WT and *tpte*$^{-/-}$ larvae. (C) Body length measurements in WT vs *tpte*$^{-/-}$ larvae, from 2 to 5 dpf. (D) WT and *tpte*$^{-/-}$ uncut fins at 5 dpf. (E) Fin lengths in uncut and cut conditions in WT vs *tpte*$^{-/-}$, from 2-5 dpf. Mean +/- SD. (F) Fin lengths in development (pink) and regeneration (purple) in *tpte*$^{-/-}$, between 2-5 dpf (individual data from E). (G) Change in fin length between d2 and d5 in *tpte*$^{-/-}$ vs WT cut. ***$p \leq 0.001$, two-tailed, unpaired, parametric t-test. (H) Top, 2dpf fin before and after cut (red dash), in *tpte*$^{-/-}$ mutants combined with transgenics labeling intracellular $Ca^{2+}$ (ubb:GCaMP6f). Bottom, Position of activated $Ca^{2+}$ wavefront relative to wound vs. time of $Ca^{2+}$ wavefront activation. Bottom inset, log-to-log scale. Cyan lines, diffusive fit in linear scale (D: diffusion coefficient), linear fit ($\alpha$: scaling exponent) in log-to-log scale (0.1s - 5s). Error, 95% CI of parameter fits. R2, goodness of fit. N=8 larvae analysed. (I) Fraction of proliferative cells vs. position from 1h to 11h uncut (dash line) and cut (full line) in *tpte*$^{-/-}$. Zero, notochord tip. Shaded areas, SD. (J) Heatmaps of position probability density of EdU+ nuclei from 3h to 11h in *tpte*$^{-/-}$. Outline, average fin boundary. Dashed line, notochord tip. (K) Comparison of cell proliferation in uncut fins between all conditions tested in this study. Scale bars: (B) 500 $\mu$m, (D) 200 $\mu$m. For A,C,E,F,G, Mean +/- SD. n: number of larvae.

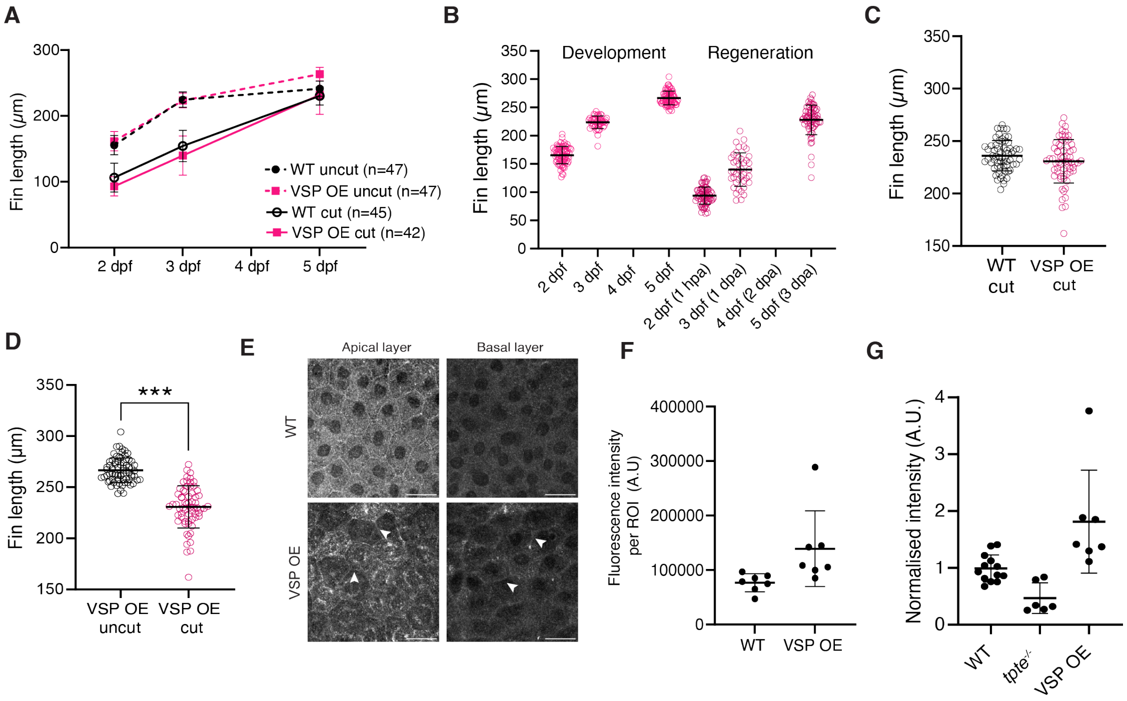


FIG. S13: **VSP overexpression induces fin developmental overgrowth and complete fin regeneration.**(A) Fin lengths in uncut and cut conditions in WT vs VSP OE, from 2-5 dpf. Mean +/- SD. (B) Fin lengths in development (pink) and regeneration (purple) in VSP OE, between 2-5 dpf (individual data from A). (C) Comparison of fin lengths at 5 dpf, in WT vs VSP OE regenerated (cut) larvae (from B). (D) Comparison of fin lengths at 5 dpf, in cut vs uncut conditions in VSP OE transgenics (from B).***$p \leq 0.001$, two-tailed, unpaired, non-parametric Mann-Whitney tests. (E) Immunostainings with anti-VSP antibody showing VSP expression in 2dpf fins per fin layer. Apical (left) and basal (right) fin epithelial cells in WT (top) and VSP OE (bottom) conditions. VSP overexpression leads to VSP enrichment in apical as well as basal epithelial layers (white arrows). Max projection of two planes. Scale bars, 20 $\mu$m. (F) Comparison of VSP immunostaining intensity between WT and VSP-OE conditions, from mean intensity projections comprising both apical and basal fin epithelial layers. n = 7 embryos for WT, 7 embryos for VSP OE. (G) Comparison of intensity of VSP immunostainings from mean intensity projections of both apical and basal fin epithelial layers normalised to mean WT intensity, in WT, *tpte*$^{-/-}$ and VSP OE conditions. n = 12 embryos for WT, 6 embryos for *tpte*$^{-/-}$ and 7 embryos for VSP OE. Mean +/- SD.

### SUPPLEMENTARY MOVIE LEGENDS

*Movie S1.* **$Ca^{2+}$ wavefront propagation in the 2dpf zebrafish caudal fin (see Fig.1C, S2A)** Movies show first 45s post-injury, induced by UV-laser ablation of the fin. Top, double transgenics labelling apical epithelial fin cells and intracellular calcium (Claudinb:Lyn-GFP;ubb:GCaMP6f); Bottom, double transgenics labelling basal epithelial fin cells and intracellular calcium (tp63:CAAX-GFP; ubb:GcaMP6f). Red line, position of laser ablation between -100 ms and 0 ms. Scale bar: 50 $\mu$m.

*Movie S2.* **$Ca^{2+}$ wavefront propagation induced by wounds of different shapes (see Fig. S3A-B)** Double transgenics labelling the apical (top, Claudinb:Lyn-GFP; ubb:GCaMP6f) or basal epithelial layer (bottom, tp63:CAAX-GFP; ubb:GCaMP6f). Left, laser ablation in a point-like injury (red triangle). Right, laser ablation in a line-shaped injury (red line). Note that all videos here were taken with a cutting setup where time-lapse imaging before and after laser ablation were executed as separate experimental blocks (Methods). Scale bar: 50 $\mu$m.

*Movie S3.* **$Ca^{2+}$ wavefront propagation upon mechanical fin injury (see Fig. S3C).** Calcium intracellular signal from ubb:GCaMP6f transgenics. Movie plays at real time speed. Scale bar: 100 $\mu$m.

*Movie S4.* **$Ca^{2+}$ wavefront propagation constructed from the minimal nonlinear model (see Fig. 2C-D).** Each rectangle represents a cell centered at position $x$. The white level of the whole cell corresponds to the calcium concentration at position $x$ and time $t$.

*Movie S5.* **$Ca^{2+}$ wavefront propagation in 2dpf zebrafish incubated in $Ca^{2+}$-free medium (see Fig. S5A).** Movies show first 15s post-injury, induced by UV-laser ablation of the fin. Top, double transgenics labelling apical epithelial fin cells and intracellular calcium (Claudinb:Lyn-GFP;ubb:GCaMP6f), compared to E3 control (left); Bottom, double transgenics labelling basal epithelial fin cells and intracellular calcium (tp63:CAAX-GFP; ubb:GcaMP6f), compared to E3 control (left). Red line, position of laser ablation between -100 ms and 0 ms. Zebrafish larvae were pre-incubated in $Ca^{2+}$-free medium (Methods). Scale bar: 50 $\mu$m.

*Movie S6.* **$Ca^{2+}$ wavefront propagation in 2dpf zebrafish with blocked gap junction pore formation (see Fig. S5B).** Movies show first 15s post-injury $Ca^{2+}$ responses, induced by UV-laser ablation of the fin, compared to control (left). Double transgenics labelling apical epithelial fin cells and intracellular calcium (Claudinb:Lyn-GFP;ubb:GCaMP6f) were used. Red line, position of laser ablation between -100 ms and 0 ms. Zebrafish larvae were injected with GAP27 peptide to inhibit gap junction pore formation (Methods). Note that this video was taken with a cutting setup where time-lapse imaging before and after laser ablation were executed as separate experimental blocks (Methods). Scale bar: 50 $\mu$m.

*Movie S7.* **$Ca^{2+}$ wavefront propagation in 2dpf zebrafish caudal fin basal epithelial layer with blocked gap junction pore formation (see Fig. S5C).** Movies show the first 15s post-injury $Ca^{2+}$ responses, induced by UV-laser ablation of the fin, compared to control condition (left). Double transgenics labelling basal epithelial fin cells and intracellular calcium (tp63:CAAX-GFP; ubb:GcaMP6f) was used. Red line, position of laser ablation between -100 ms and 0 ms. Zebrafish larvae were injected with GAP27 to inhibit gap junction pore formation (Methods). Scale bar: 50 $\mu$m.

*Movie S8.* **$Ca^{2+}$ wavefront propagation in KCl-supplemented media (see Fig. 2E).** Injury $Ca^{2+}$ responses post UV-laser ablation of the 2dpf fin, between WT (Top, E3 medium incubated) and KCl-supplemented (Bottom, 135 mM KCl supplemented-E3 medium). Double transgenics labelling intracellular $Ca^{2+}$ and apical epithelial layer (Claudinb:Lyn-GFP; ubb:GCaMP6f) were injured with a UV-laser between -100 ms and 0 ms. Time-lapse images with an extended field-of-view were taken starting from 10 s post-injury. Scale bar: 100 $\mu$m.

*Movie S9.* **$Ca^{2+}$ wavefront propagation in $K^{+}$ flux perturbations (see Fig. S6).** Injury $Ca^{2+}$ responses within the first 45 s post UV-laser ablation of the 2dpf fin, between WT (top left), KCl-supplement (top right), *lof*$^{-/-}$ (bottom left) and FK506-treatment (bottom right). Note that WT (top left) shows the same dataset as in Movie S1 and Fig. 1C. Double transgenics labelling intracellular calcium andapical epithelial layer (Claudinb:Lyn-GFP; ubb:GCaMP6f) were used in all conditions. Red line, position of laser ablation between -100 ms and 0 ms. Scale bar: 50 $\mu$m.

### MATERIALS AND METHODS

### EXPERIMENTAL PROCEDURES

#### 1. Ethics Statement

This study followed European Union directives (2010/63/EU) and German law, with license # TVV52/2021 -*'Generierung von Zebrafischlinien zur Untersuchung der Größe und Form von Organen und Organellen'*. Genetic engineering work was carried out in an S1 area (MPI-CBG, S1-Labore 4., Az.: 54-8451/103, project leader Rita Mateus), following guidelines according to Section 21, Paragraph 1 of the German Genetic Engineering Act, and within projects 01-03 from the Mateus laboratory, according to Section 28 of the German Genetic Engineering Safety Ordinance (GenTSV).

#### 2. Zebrafish lines and maintenance

All zebrafish (*Danio rerio*) lines were maintained in a recirculating system with a 14 h/day, 10 h/night cycle at 28°C. Crosses were performed with 3- to 12-month-old adults. Embryos were kept in E3 zebrafish embryo medium [1] at 28.5°C until the desired developmental stage was reached. We used the following transgenics either in WT AB or $longfin^{t2/t2}$ (i.e. $lof^{-/-}$) [2] backgrounds: $Tg(\text{-}8.0cldnb\text{:}lynGFP)^{zf106}$ [3] (i.e. claudinb:Lyn-GFP) for labelling the fin's apical epithelial cell layer (periderm); $Tg(tp63\text{:}CAAX\text{-}GFP)^{mdi2013Tg}$ [4] for labelling the fin's basal epithelial cell layer; and $Tg(actb2\text{:}LY\text{-}tdTomato)^{zf2254Tg}$ (i.e. b-act:lyn-tdTomato) to label all membranes [5].

#### 3. $tpte^{cbg22}$ mutant

$tpte^{cbg22}$ CRISPR mutants were generated as described previously [6]. Briefly, trans-activating crispr RNAs (crRNAs) were designed for specific loci of the *tpte* gene (Ensembl ID: ENSDARG00000056985) using pre-designed crRNAs (IDT). Several crRNAs were tested for ribonucleoprotein (RNP) mutagenesis and were chosen considering where the start codon is located, as well as high on-target and low off-target scores. The most efficient crRNA tested was located on exon 5 of the zebrafish tpte canonical transcript, with target sequence 5'-

CACGGAGCAATACATCGATG-3' (Dr.Cas9.TPTE.1.AC, IDT). The crRNA was annealed with an equal molar amount of tracrRNA (#1072533, IDT) and diluted to 57 $\mu$M in Duplex buffer (#11-01-03-01, IDT), generating the single guide RNA (sgRNA). The sgRNA was mixed with the Cas9 protein (Alt-R S.p. Cas9 Nuclease V3; 1081058, IDT; 61 $\mu$M stock) in equimolar amounts, generating a 28.5 $\mu$M RNP solution. To improve mutagenesis efficiency, the mix was kept overnight at -20°C before injections the following day. One-cell stage *Tg(ubb:GCaMP6f)*$^{m1299}$ embryos were injected with 1 nl of the RNP mix and grown to adulthood. Founder fish containing frameshift mutations were identified by genotyping the resulting F1 progeny, resulting from outcrosses with WT AB fish. The mutant fish showed a 4bp deletion in exon 5 (Fig 4C).

### 4. Transgenic lines generation

All transgenesis was performed using the Tol2 transposon system and Gateway cloning [7, 8] technology. For *Tg(Hsp70:CiDr-VSP-L223f-mCherry)*$^{cbg23Tg}$ generation, we used the CiDr-VSP-L223F-mCherry pcDNA3.1 plasmid [9] (addgene #140892) as a template for subcloning. The CiDr-VSP-L223F-mCherry fragment was amplified using primers containing attB1 and attB2 sites: FWD 5'-GGGGACAAGTTTGTACAAAAAAGCAGGCTTAatggagggattcgacggttca-3'; REV 5'-GGGGACCACTTTGTACAAGAAAGCTGGGTTttacttgtacagctcgtccatgcc-3'. The 2494bp amplified product was purified using a PCR clean-up kit (Promega, #A9281), and recombined with pDONOR221 (#208, Tol2 Kit [8]) in a BP reaction (BP Clonase II enzyme mix; Invitrogen, #11789020). The resulting middle entry clone, pME_CiDr-VSP L223F-mCherry, was then recombined in a LR reaction (LR Clonase II Plus enzyme; Invitrogen, #12538120) using p5E-hsp70 (#222, Tol2 Kit), pDEST-Tol2pA_myl7:EGFP (R4-R3) (#395, Tol2 Kit) and p3E-polyA (#302, Tol2 Kit), to generate the final construct, T2_Hsp70:CiDr-VSP-L223F-mCherry_pA_myl7:GFP. One-cell stage WT AB embryos were injected with 25 pg of the final construct, 25 pg of Tol2 transposase mRNA and 10% phenol Red. At 2 dpf, injected embryos were screened for GFP expression in the heart (driven by the myl7 selection marker) using an Olympus SZX16 fluorescence microscope, and subsequently grown to adulthood. Founder fish were identified by screening F1 progeny for selection marker fluorescence, resulting from outcrosses with WT AB fish.

For the *Tg(ubb:GCaMP6f)*$^{m1299}$ line, an attB1 primer (5'-GGGGACAAGTTTGTACAAAAAAGCAGGCTGAACCGTCAGATCCGCTAG-3') including Kozak sequence was used to amplify GCaMP6f [10] (Addgene plasmid #40755 pGP-CMV-GCaMP6f). The ubb promoter (3.5 kb [11], Addgene plasmid #27320) was inserted into the pENTR5' plasmid. pENTR5' (ubb), pME (GCaMP6f), and pENTR3' (polyA) sequences were then cloned into the pDestTol2pA2 plasmid via an LR clonase reaction (Thermo Fisher Scientific Gateway LR Clonase Plus enzyme; #12538120). 25 ng/$\mu$l plasmid DNA and 50 ng/$\mu$l Tol2 transposase mRNA were co-injected into single-cell stage embryos (*nacre*$^{+/-}$ background). F2 or later generations in wildtype AB background were used for this study's data acquisition.

**5. Genotyping**

Genomic DNA from individual larvae or clipped tail fins from adult zebrafish were extracted using the Kapa Express Extract kit (Kapa Biosystems) according to the manufacturer's protocol. This was followed by performing PCR with KAPA2G Robust HotStart ReadyMix (Kapa Biosystems). For *tpte*$^{cbg22}$ genotyping, the following primers were used: FWD, 5'-GTTTTCGGTGAGTGGCATAC-3'; REV, 5'-GTTACTGTTAGTCTTACGAGC-3'. Resulting PCR from either mutant products were sent for sequencing and sequences analyzed using Snapgene (v7.1).

For *longfin*$^{t2}$ genotyping, the following primers were used based on [12]: *wt_breakp_fwd*, 5'-cgttgaatcaccgttgaaatgcc-3'; *lof_breakp_fwd*, 5'-ggccttgttaagctcaagtg-3'; and *breakp_rev*, 5'-ggtttcgatatgtggcagatttaagg-3'. Primer pairs were used in combination in two concomitant PCRs (alternating only the FWD primer), resulting in PCR products of 522 bp (*wt_breakp_fwd* and *breakp_rev*, amplifying a sequence of WT gene) and 526 bp (*lof_breakp_fwd* and *breakp_rev*, amplifying a sequence of *lof* gene) for heterozygous fish, and only the 526 bp product for homozygous fish. Given that *lof* is a dominant mutation [2], fish with fins that present wildtype dimensions were not genotyped. Genotyping was performed to identify heterozygous and homozygous fish, despite phenotypic differences in fin lobe size.

### 6. Voltron2 Cloning, mRNA synthesis and embryo microinjection

The construct pCS2+-Voltron2 was generated by subcloning from pGP-pcDNA3.1 Puro-CAG-Voltron2 [13] (Addgene #172909) with Gateway technology [7, 8]. Briefly, the Voltron2 fragment was amplified using primers containing attB1 and attB2 sites: FWD 5'-GGGGACAAGTTTGTACAAAAAAGCAGGCTTAatggctgacgtggaaaccg-3'; REV 5'-GGGACCACTTTGTACAAGAAAGCTGGGTTttacacctcgttctcgtagcagaac-3'. The 1720 bp amplified product was purified using a PCR clean-up kit (Promega, #A9281), and recombined with pDONOR221 (plasmid #208, Tol2 Kit [8] in a BP reaction (BP Clonase II enzyme mix; Invitrogen, 11789020). The generated middle entry clone, pME_Voltron2, was then recombined in a LR reaction using pCSDEST (plasmid #201, Tol2 Kit) (LR Clonase II Plus enzyme; Invitrogen, 12538120) to generate the pCS2+-Voltron2. To make Voltron2 mRNA, the pCS2+-Voltron2 vector was linearised with NotI (NEB) and transcribed using the SP6 mMessage Machine Kit (#AM1340, Invitrogen) according to manufacturer's instructions. The same procedure was applied for Tol2 transposase mRNA synthesis, generated from a pCS2FA-transposase plasmid. mRNAs were aliquoted and stored at -70 °C until use. One-cell stage wildtype AB or claudinb:lyn-GFP trangenic embryos were injected using standard procedures with 150-200 pg of Voltron2 mRNA. Embryos were left to develop at 28.5 °C until the desired stage. A PV-820 Pico-injector (World Precision Instruments) and a Narashige micromanipulator were used for microinjection.

### 7. GAP27 peptide heart injections

To inhibit pore formation of gap junctions, 0.5 mM Gap27 peptide (700 pg per embryo) (Tocris, #1476) was injected in the heart of 2 dpf larvae as previously described [14]. Control embryos were injected with a mix of water and phenol red, without GAP27 peptide. Briefly, 1mm borosilicat glass needles (Harvard Apparatus, Premium thin Wall Borosilicate Capillary Glass, OD 1.0 mm, ID 0.78 mm, #640778) were pulled with a TCF Sutter "9" to the inner diameter of 15 $\mu$m, with a bevel at 30°and a spike, to facilitate piercing into the zebrafish skin. Embryos were anaesthetized with 20 mg/ml of tricaine [15] and were placed against a glass slide in a plastic petri dish with the head facing the slide, the needle was positioned at a 30-45°angle with respect to the heart, and it was introduced in the atrium without piercing the yolk. Embryos were imaged 5h after injection due to known Connexin turnover [16, 17]. A PV-820 Pico-injector (World Precision Instruments) and a Narashige micromanipulator were used for microinjection.

### 8. Chemical treatments

Prior to all chemical treatments, embryos were manually dechorionated.

#### *1. Voltron2 labelling with JF552*

Embryos injected with Voltron2 mRNA were incubated with 500nM of Janelia Fluor Dye 552-HaloTag ligands in E3 with 0.5% dimethylsulfoxide (DMSO) at 36 hpf, for 12h. The dye was then washed twice for 10 min with 4 mL of E3 with 0.5% DMSO. Embryos were screened for fluorescence (Olympus SZX2 with a X-Cite Series 120PC Q LED lamp, ET 630/75 bandpass) before mounted for live imaging in 0.5% low-melt agarose (E3).

#### *2. $Ca^{2+}$ free E3 medium*

$Ca^{2+}$ free 1x E3 medium (dissolved from 60x solution: 34.8 g NaCl + 1.6 g KCl + 9.78 g $MgCl_2 \cdot 6H2O$ in 2 L $H_2O$) was prepared and used to incubate embryos for 5h prior to imaging. Up to 40 embryos were incubated per 15 mL of medium. Embryos were mounted for live imaging in 0.5% low-melt agarose ($Ca^{2+}$ free E3).

#### *3. KCl supplement*

KCl supplemented medium consists of concentrated KCl in E3 medium (135 mM, ∼282 mOsm/L, considered isotonic to the fin's interstitial fluid [18]) (5.032g KCl dissolved in 500 mL E3, Supelco, CAS #7447-40-7). Two treatment conditions were employed, where no harmful side effects were noted in embryos: a) overnight incubations and b) 1h incubations. In the first case, embryos were incubated in KCl-supplemented medium at 36 hpf, for 12-18 h until imaging of $Ca^{2+}$ wave (Fig. 2G-H). Up to 40 embryos were incubated per 15 mL of KCl-supplemented medium. After the incubation, embryos were mounted for live imaging in 0.5% low-melt agarose (made with KCl-supplemented E3 medium). In this overnight KCl treatment condition (without cutting), we observed a longer caudal fin at 5 dpf (Fig. S11F) that suggests that an increase in cell proliferation took place. The second KCl treatment (1h) was performed to enable the capturing of cell proliferation events. In the case of monitoring proliferation induced by injury-independent depolarisation, uncut larvae were incubated for 1h in 135mM KCl (with EdU, see below). In regeneration conditions (cut fins), KCl was incubated 1h prior to amputation and washed 1h after amputation (with EdU, see below). We also examined the injury-induced $Ca^{2+}$ wave response under this 1h KCl treatment condition. To do this, we mounted 48 hpf embryos in 0.5% low-melt agarose (E3 medium), then perfused in the dish KCl-supplemented medium at 1h before laser microdissection and live imaging. In control conditions, only E3 medium was perfused. Intriguingly, in this condition, we found that $Ca^{2+}$ wave still stops at $> 300$ $\mu$m away from the cut (Fig. S11G-H), agreeing with the 3-fold increase displayed in the overnight KCl treatment condition. While osmotic effects may occur after the injury electric responses (90s to min timescale, [18]), the KCl treatment(s) applied here did not disrupt regeneration and promoted fin growth (Fig. 3K-N, S11).

#### *4. FK506*

Embryos were incubated with 10 $\mu$M FK506 [19] (Sigma #F4679) at 36 hpf, for 12-18h prior to live imaging. Up to 40 embryos were incubated per 15 mL of FK506-supplemented medium. After the overnight incubation, embryos were mounted for live imaging in 0.5% low-melt agarose (FK506-supplemented E3 medium). For regeneration and proliferation assays, FK506 was washed at 1 hpa.

#### 9. Spinning disk live imaging and UV-Laser microdissection

Prior to imaging, embryos were anesthetized with 0.1% Tricaine (MS-222, Sigma) dissolved in E3 medium. Anesthetized embryos were mounted in 0.5% low-melt agarose (Sigma) containing 0.1% Tricaine in glass bottom dishes (CellView). Live imaging was performed at 28.5°C (stage-top incubator) using a spinning disk confocal microscope equipped with a Yokogawa CSU-X1 scan head (25x water immersion objective, 0.8 NA; paired with an objective heater also at 28.5°C) connected to a Zeiss camera (AxioCam 705 mono). Images were acquired using the Zeiss ZEN (3.2 blue) software. Caudal fin microdissection was performed via 355 nm laser ablation (1 kHz rep rate, max 42 $\mu$J pulse power, 1 ns pulse length) implemented through a RAPP Optoelectronic module that is scanner-based coupled to the microscope. The RAPP module runs on the SysCon2 software (2.3.1), also coupled to the ZEN software through the Image Transfer function. For both GCaMP6f (488 nm excitation, BP 535/30) and Voltron2:JF552 (561 nm excitation, BP 629/62) imaging, continuous imaging (100 ms per frame) in a single z-plane was performed while ablation was triggered at a pre-specified timeframe (with a long pass beamsplitter T387LP in place). The scanning speed and position of the ablation laser are kept invariant across embryos, such that the microdissection takes place at $\sim$50 $\mu$m away from the tip of the fin and the triggered ablation laser traverses the tissue in 100 ms. The ablation laser power was estimated and adjusted per embryo to minimize positional shifts and cavity bubbles (due to excessive heating) while generating a line wound. While most UV-laser ablation experiments were performed using the continuous-imaging setting described above (Fig. 1,2,S2,S4,S5A,S5C,S6; Movies S1,S5,S7,S8,S9), the experiments in Fig. S3, Fig. S5B and Movies S2,S6 followed a UV-laser ablation regime where time-lapse imaging before and after laser ablation were executed as two separate experimental blocks. In this setting, the first frame of the second continuous imaging block (130 ms per frame) was taken as 0s.

#### 10. Embryo heat-shocks

Heat-shocks were performed at 37°C for 1 hour in a water-bath, where up to 100 embryos were placed into a petri-dish containing 30 mL of E3. For VSP overexpression experiments with *Tg(Hsp70:CiDr-VSP-L223f-mCherry*$^{cbg23Tg}$*)*, two heat-shocks were performed: the first at 31 hpf, the second at 42 hpf. Embryos were sorted for fluorescence and/or amputated after 6 hours from the last heat-shock (48 hpf).

#### 11. Mechanical amputation and injury

Mechanical fin fold amputations were performed at 48 hpf, as described previously [20, 21]. Prior to amputation, embryos were anesthetized with 0.1% Tricaine (MS-222, Sigma) dissolved in E3 medium.

*1. Fin fold regeneration assay*

To study the effects of all perturbations (drugs, mutants, overexpression) during regeneration, fin folds were amputated with a 15 mm surgical razor blade (Braun) at a position posterior to the notochord. After, embryos were returned to E3 medium and incubated at 28.5°C until desired developmental stages, when they were further processed for live imaging, immunostaining or proliferation assays. When live-imaged, uncut and cut larvae were placed in a 96 well plate (CellView) in E3 with tricaine and imaged in brightfield mode with an automated live cell confocal microscope CellVoyager CV7000 (Yokogawa) with a 4x/0.16 air objective at 2, 3 and 5 dpf.

*2. Stereoscope $Ca^{2+}$ imaging following fin injury*

To live-capture $Ca^{2+}$ signals following mechanically-induced injury, anesthetized fish were placed in a plastic petri dish (Greiner Bio-One) and fin folds were wounded with a handheld plastic pipette. Videos were taken with an iPhone 13 connected to the eyepiece of a stereoscope (Olympus SZX2 with a X-Cite Series 120PC Q LED lamp, ET 510 bandpass) by a smartphone adapter (Dörr SA-1).

#### 12. Proliferation assay

To label proliferating cells, the EdU Click-iT-Alexa 647 fluorophore kit (Thermofisher Scientific #C10340) was used. Briefly, larvae from different developmental or amputation timepoints were incubated at 4°C in E3 medium with 500 $\mu$M of EdU in 10% DMSO for 1 hour. Embryos were then washed three times with E3, and fixed overnight in 4% PFA at 4°C. After wholemount immunostaining for p63 to label basal epithelial cells, EdU detection was performed as per the manufacturer's protocol. Embryos were stored in 80% Glycerol, 2% DABCO (Sigma) at 4°C, protected from light until imaging.

#### 13. Immunofluorescence

Whole-mount immunostainings were performed as previously described [21]. The primary antibodies used were: rabbit anti-p63 1:200 (Abcam, #ab97865), mouse anti-VSP 1:100

(BIOZOL, #75-485), rabbit anti-mCherry living colours (Clonetech, #632496), mouse anti-ZO-1 (#339100, Thermofisher Scientific), mouse anti-E-cadherin (#610181, BD Bioscience) and rabbit anti-Cx43 (#3512S, Cell Signaling). The secondary antibodies used were: alexa fluor 488 goat anti-rabbit (#A11034, Life technologies), alexa fluor 488 goat anti-mouse IgG2a (#A21131, Thermofisher Scientific), alexa fluor 488 goat anti-mouse (#A11001, Life technologies), alexa fluor 594 goat anti-mouse IgG2a (#A21135, Thermofisher Scientific) (all used at 1:500 dilution). DAPI (#D9564, Sigma) was incubated with the secondary antibodies at 1:1000 dilution. Immunostainings were repeated at least 2 times with different biological replicates, per marker and condition. Embryos were mounted for imaging in 0.5% low-melting agarose (A9414, Sigma) diluted in E3 or 80% Glycerol, 2% DABCO (Sigma) diluted in PBS. Images were acquired with an inverted Zeiss LSM 880 AiryScan point-laser scanning confocal microscope equipped with a C-Apochromat 40x/1.2 water immersion objective (Zeiss) or a Zeiss Cell Discoverer with a Plan-Apochromat 50x/1.2 water immersion objective (Zeiss). For imaging Connexin43 stainings, we used a Zeiss Lattice Lightsheet 7 equipped with a 13.3x/0.4 water immersion illumination objective and a 44.83x/1.0 water immersion detection objective, allowing isometric voxel resolution. Images were acquired using the software Zen Blue (v3.10.103) for the light sheet, and software Zen 2011 Black edition and Zen Blue v. 3.6.095.09000 for laser scanning confocal systems.

#### 14. Transmission electron microscopy (TEM)

2 dpf wildtype AB larvae were fixed in 2% Paraformaldehyde, 2% Glutaraldehyde, 50 mM Hepes (pH 7.2) at room temperature for 1 hr, then transferred to 4°C overnight. Then, the fixative was washed 3x5 min with 50 mM Hepes and rinsed with water, prior to contrasting with reduced osmium tetroxide (1% $OsO_4$ (EMS), 1.5% potassium ferrocyanide (Sigma)) in water for 1h at room temperature. Samples were then washed with ultrapure water at least three times between each staining/contrasting step. Staining was enhanced by incubation with 0.2% tannic acid (EMS) for 15 min at room temperature. A further contrasting step with 0.5% uranyl acetate (EMS) in water followed for 1h at RT in the dark. The tissue was further washed with water and then gradually dehydrated with increasing concentrations of ethanol (starting at 30% ethanol and finishing with 3x 100% ethanol incubation for 15, 15 and 30 min), followed by a stepwise infiltration with epoxy resin:ethanol 1:2, 1:1, 2:1

(EMBed812, EMS). A final infiltration step in pure resin was performed at least overnight. The resin was finally cured at 60°C for at least 24 h. For conventional TEM analysis, 70 nm–sections were cut on a Leica UCT ultramicrotome (Leica Microsystems, Wetzlar, Germany) onto formvar-coated slot grids and post-stained with 2% aqueous uranyl acetate and lead citrate (EMS). Sections were collected at the notochord level (∼160$\mu$m from fin tip, Fig. S1C) and at 35µm from the fin tip (Fig. 1B). Images were taken on a Tecnai Biotwin T12 electron microscope (Philips/Thermofisher Scientific) with a F416 CCD camera (TVIPS).

### QUANTITATIVE PROCEDURES

#### 1. Epithelial cell segmentation and registration

To obtain cell segmentation, raw image stacks collected from Zen Blue software were imported via ImageJ (v2.14.0) and processed via the "cyto" model in Cellpose [22] (v2.2.3, auto-calibrated cell diameter per image stack). Segmented masks from Cellpose were then imported to custom MATLAB (R2022b) scripts for following single cell registration and analyses. To account for the tissue contraction that becomes prominent after ∼3 s after laser microdissection (Fig. 1J, Fig. S4), two different cell registration strategies were employed for GCaMP6f and Voltron2:JF552 signals. For apical GCaMP6f signals that were evaluated for up to 45 s after microdissection, we registered a list of moving cell positions by first segmenting the double transgenics (ubb:GCaMP6f;claudinb:Lyn-GFP) at every time frame, then tracking the center-of-mass across all time frames (Hungarian-based particle linker [23], distance cutoff for frame-to-frame linking set to 10 pixels = 2.76 $\mu m$, no particle disappearance gap allowed). A round of manual correction was then performed (custom MATLAB script) to validate the tracked cells and remove artifacts. For apical and basal Voltron2:JF552 signals that were evaluated for only up to 3 s after microdissection, we registered a list of static cell positions by segmenting the membrane reference image (see next section "Voltron2:JF552 membrane signal quantification") at the time frame 100 ms before laser microdissection was applied. In this case, a round of manual correction was performed in the GUI interface of Cellpose to remove artifacts. We note that for basal GCaMP6f signals, a list of static cell positions were also used because of the limited accuracy of Cellpose's "cyto" model for segmenting double transgenic ubb:GCaMP6f and tp63:CAAX-GFP, in particular when GCaMP6f has a high cytosolic intensity. In this case, we evaluated the single cell GCaMP6f intensity for up to 10 s after microdissection (Fig. S2C). We also performed cell-independent wavefront evaluation of GCaMP6f change of intensity to ensure a reliable comparison with apical GCamp6f signals (Fig. S2B).

#### 2. GCaMP6f cytosolic signal quantification

To quantify single cell GCaMP6f dynamics in layer-specific datasets (from double transgenics ubb:GCaMP6f;claudinb:Lyn-GFP or ubb:GCaMP6f;tp63:CAAX-GFP), a list of cell mass centers and boundaries was first registered via segmentation (see the section before). For each cell, the cytosolic GCaMP6f intensity was then extracted as the mean intensity of pixels enclosed by a proportionally-contracted cell boundary (80% the distance to the center of mass $r_c$ compared to the segmented boundary $r_s$, $r_c yto = 0.8 * (r_s - r_c) + r_c$). In order to compare $Ca^{2+}$ activation dynamics across embryos that could have different GCaMP6f insertion sites, the cellular change of GCaMP6f intensity (as compared to the GCaMP6f intensity before laser microdissection, $\Delta F = F - F_{uncut}$) was first normalized by the maximum intensity change across all cells ($\Delta F_{max}$), a quantity specific to each embryo. The single cell $Ca^{2+}$ activation time was then determined by evaluating the time when this normalized cellular change of GCaMP6f intensity ($\Delta F/\Delta F_{max}$) surpasses a threshold (fixed to 0.1 throughout all analyses) for the first time. Based on the sub-second sharp increase of GCaMP6f intensity for cells close to the cut (Fig. 2C), we also performed a 10-point linear interpolation (10 ms per interpolated time point, compared to raw intensity time-lapse at 100 ms per frame) between consecutive time points to extract an accurate estimate of when $\Delta F/\Delta F_{max}$ surpasses 0.1. Fitting the single cell distance from cut (defined as its mass center from the line instructed for laser microdissection) as a function of the activation time (non-linear least square method, MATLAB) then returns the fitted $Ca^{2+}$ wavefront activation dynamics (Fig. 1F). In $lof^{-/-}$ and FK506-treated embryos, a prominent fraction of cells spatially disconnected from the $Ca^{2+}$ wavefront showed rapid $Ca^{2+}$ activation within the initial seconds post injury (i.e. Calcium flashes, Fig. 2I-J), possibly resulting from increased expression of *kcnh2a* [24], or increased $K^+$ leak through *kcnk5b* [25–27] in the epithelium. These cells were excluded (manual filter, custom MATLAB script) in fitting of the $Ca^{2+}$ wavefront dynamics (Fig. 2J, Fig. S6). On the other hand, $Ca^{2+}$ flashes were determined by detecting local peaks ("islocalmax" function, MATLAB) in the normalized single cell change of GCaMP6f intensity ($\Delta F/\Delta F_{max}$) that is not identified to be a first-time $Ca^{2+}$ wavefront activation from above. A threshold of prominence (defined as the maximum height this local peak displays relative to neighboring valleys, threshold fixed to 0.15 throughout all analyses) was used to keep only those prominent local peaks. The detected peaks were finally passed through a round of manual proofread to ensure the absence of artifacts (Fig. 1F, Fig. 2H, Fig. 2J).

In addition to layer-specific single cell GCaMP6f quantification, an additional quantification pipeline was developed to evaluate GCaMP6f intensity changes in datasets without epithelial membrane markers. This was performed for ubb:GGCaMP6f in the background of $tpte^{-/-}$ (in comparison to WT, Fig. S12H). In this pipeline, GCaMP6f pixel intensity changes ($\Delta F$) were also normalized by the maximum intensity change of all pixels ($\Delta F_{max}$) for each embryo. Instead of evaluating the activation time per single cell, pixels enclosed by the boundary of the caudal fin (the boundary was created manually and excludes evident fibroblasts or notochord composition) were binned ($20\mu m$ width, based on segmented cell size, Fig. S8G) according to their distances from cut, and activation time was defined as when the average normalized intensity change in a bin surpasses the same threshold ($\bar{\Delta F}/\Delta F_{max} > 0.1$) for the first time. The binned distance plotted vs. the bin activation time then constitutes a simplified evaluation of the concomitant wavefront propagation dynamics for both apical and basal epithelial layers (Fig. S12H). Lastly, for datasets with epithelial membrane markers, we also employed the membrane marker-independent approach to evaluate the position and instantaneous velocity of the $Ca^{2+}$ wavefront. This was done by plotting the normalized change of GCaMP6f intensity as a function of distance (no binning; distance is in unit of pixels) for every time frame, and annotating the position where this normalized value drops to 0.1 as the instantaneous wavefront position (Fig. S2B). Evaluating the instantaneous wavefront velocity (defined as the wavefront position after 1s=10 frames minus the current position) then determines the time when the wavefront velocity drops to 0 (Fig. 1D). This time (~5s for both apical and basal layers) was thus used as the time range for fitting the $Ca^{2+}$ wavefront activation dynamics. Custom MATLAB scripts for GCaMP6f signal analyses as summarized in this section are available at [28].

#### 3. Voltron2:JF552 membrane signal quantification

To quantify single cell membrane Voltron2:JF552 signals in layer-specific manner (Voltron2 mRNA-injected into claudinb:Lyn-GFP or tp63:CAAX-GFP, then embryos stained with JF552), we first obtained a list of static cell boundaries via segmentation of the membrane reference image (see the section "epithelial cell segmentation and registration"). Here, we note that because of the variability of Voltron2 expression and JF552 staining at the epithelial layers, the membrane reference image is either the before-cut claudinb:Lyn-GFP or tp63:CAAX-GFP (when Voltron2:JF552 primarily appears at the epithelial layer labelled by transgenic membrane marker); or the before-cut Voltron2:JF552 image (when Voltron2:JF552 primarily appears at the epithelial layer not labelled by transgenic membrane marker). In the latter case, we manually exclude cells whose membranes show both the transgenic and the Voltron2:JF552 signals (GUI interface of Cellpose) in order to ensure all registered cells come from the same epithelial layer. Next, these segmented cell boundaries were used to determine position of the membrane pixels. This was defined by extracting all pixels enclosed by a sliding square window with the radius of 3 pixels=0.828µm on the cell boundary. Here, the effective "membrane thickness" (6 pixels=1.656$\mu m$) was estimated by plotting the Tg(b-act:lyn-tdTomato) membrane marker (same imaging setup as Voltron2:JF552) intensity vs. pixel distance from membrane and extracting the half-max width of the intensity distribution. Changes of Voltron2:JF552 intensities (compared to the frame before cut, $\Delta F = F - F_{uncut}$) across all extracted membrane pixels were then binned according to their distance from cut (2 $\mu m$ width) to evaluate the raw drop of membrane Voltron2:JF552 intensities in comparison to claudinb:Lyn-GFP or tp63:CAAX-GFP membrane marker controls (Fig. 1H).

To obtain cell-averaged membrane signals, raw Voltron2:JF552 intensities from the membrane pixels from the boundary of the same cell were then grouped and averaged. The resulting cellular Voltron2:JF552 time-lapses were then passed through a filter that excludes cells with a large fluctuation even before laser microdissection onsets (here taken as the range of cellular intensity is smaller than 8% of the mean of cellular intensity before-cut) (Fig. 1I). The fractional drop of cell-averaged Voltron2:JF552 signals after laser microdissection ($\Delta F/F_{uncut}$) was subsequently determined (Fig. 1J). Finally, noting that laser microdissection of apical membrane control (claudinb:Lyn-GFP) brings about a level of membrane intensity changes (likely due to the elastic after-cut tissue movement that is more prominent in apical layers), we employed a noise floor method to fit only the cells that exhibited Voltron2:JF552 signal changes that is significant compared to membrane controls. The noise floor was constructed by binning the cell-averaged membrane signals according to their center-of-mass distance from cut (2 $\mu m$ width) for the microdissected membrane controls. Then, only Voltron2:JF552 cells within this binned distance that exhibited > two-fold change compared to this noise floor were kept. The fractional drops of these Voltron2:JF552 cells were then fitted as a function of single cell distance from cut (nonlinear least square method, MATLAB). Custom MATLAB scripts for Voltron2:JF552 signal analyses are available at [29].

#### 4. Fin and body length quantifications

Fin length (Fig. 3A) and body length were manually measured from acquired whole-body brightfield images with the line tool in ImageJ[30] v. 1.54f. Caudal fin length was measured from the end of the notochord to the tip of the fin (Fig. 3A), while body length was measured from the mouth to the tip of the caudal fin, along the notochord. Data was plotted and statistically analysed using GraphPad Prism v. 10.4.1. Before choosing the comparison test, the data was tested for normality (Gaussian distribution) using the Kolmogorov Smirnov test. In case of a normal distribution, unpaired, parametric, two tailed t-tests were used. In case of non-normal distribution, unpaired, non parametric, two tailed Mann Whitney test were used.

#### 5. Fin epithelial layer thickness and interstitial space quantification

Quantification of the thickness of the apical and basal fin epithelia were performed on images acquired at the Lattice light sheet, which allows for isotropic resolution in XYZ. Uncut caudal fins of fixed double transgenics (tp63:CAAX-GFP, b-act:lyn-tdTomato) embryos were imaged and the cumulative thickness of apical and basal epithelial layers was measured with the Fiji line tool from XZ projections at different distances from the fin tip (S1C,D).
For measurements of the fin's interstitial space, concerning area to perimeter ratio, the area between the dorsal and ventral epithelial layers was traced form the same uncut fins using

the polygon selection tool, and measurements of the area and perimeter were extracted. Measurements were performed at the usual injury position (50 $\mu$m from the tip) used for all experiments in Figure 1 and 2, and at various distances from that position (S1C,E).

#### 6. Proliferation quantification

Quantification of the proliferation patterns and levels was performed using z-stacks of larval finfolds encompassing the full organ volumes acquired at different timepoints. To quantify the percentage of proliferative cells over total cell number over time, Imaris v.10.1.1 spot detection was used to count EdU+, p63+ and DAPI+ cells, in uncut and cut fins of various developmental and regenerative times. A spot radius of $4\mu m$ was used for DAPI, and $6\mu m$ was used for EdU and p63. The Imaris machine learning algorithm was then used to differentiate nuclei in the finfold from nuclei in the notochord, which were then used to spatially align the fins but excluded from the analysis. To quantify the spatial distribution of proliferative cells in larval finfolds, the XY coordinates of each detected spot were extracted from IMARIS. A custom made Python3 script (v 0.0.1, available here: https://zenodo.org/records/15019090)[31] was generated to plot the ratio of proliferative cells (EdU+) over total cells according to their position in the fin, as a Kernel Density estimation. To align all fins, first the quantiles of the distribution of XY position of all spots were computed and used to re-orient all fins with the cut or tip of the fin to the right. Then, the PCA of the XY positions of cells in the notochord was computed, and the first component was taken to align the notochord horizontally. Bins of $20\mu m$ were used when computing the EdU/DAPI ratio, as the average cell diameter of 48 hpf fin epithelial cells is $18\mu m$ (Fig. S8G). To obtain heatmaps of the spatial distribution of proliferative cells, the fins from each group were first aligned and centered with respect to the tip of the notochord, and their datasets were merged. Then, the pixel of unit value representing the centroid of the cell (obtained from the Imaris spot detection) was plotted and a gaussian blur of $10\mu m$ (sigma, corresponding to the cell radius) was applied to each spot to simulate a point spread function. Finally, the values were normalised by the sum of the image matrix to obtain a position probability density. The values of the density were then mapped to a colormap upon plotting. To keep good resolution in the finfold and avoid cells in the notochord, the value of density was kept in a range between zero and a common clipping value of cmap.

#### 7. VSP intensity quantification

To quantify the expression of VSP in WT, *tpte*$^{-/-}$ and VSP OE, a square ROI of 96.93 $\mu m^2$ was drawn between the end of the notochord and the fin tip, and mean fluorescence intensity was measured in Fiji. All values of intensity were then normalised to the average intensity of VSP staining in WT. 2 replicate experiments were performed per condition. Data was plotted and statistically analysed in GraphPad Prism 10.4.1.

### SUPPLEMENTARY THEORY NOTES

### I. MULTI-COMPARTMENT MODEL OF ION TRANSPORT

#### 1. Model and geometry

The fin of the zebrafish larva is made of two layers of epithelial cells: apical and basal (not to be confused with the apical and basal sides of a single epithelial cell), see Fig. 1A. The apical surface of the apical epithelial cells is in contact with the external medium (E3, $\simeq$ 10 mOsm) of low osmolarity, and the basal surface of the apical epithelial cells is in contract with the interstitial fluid whose composition is of high osmolarity ($\simeq$ 270 mOsm, with a composition similar to Ringer solution). The basal epithelial cells are only in contact with the fin's interstitial fluid.
The main ions in the interstitial fluid are sodium $Na^+$, potassium $K^+$ and chloride $Cl^-$. Since the concentration of $K^+$ in the external medium and the interstitial fluid is small compared to $Na^+$, we only consider the dynamics of $Na^+$ and $Cl^-$ in the model.
Apical epithelial cells express tight junctions that compartmentalize the interstitial fluid from the external medium. The cytoplasm of apical epithelial cells is also connected through gap junctions, enabling direct cell-cell connections. Basal epithelial cells do not express tight or gap junctions. In the following sections, we therefore describe apical epithelial cells as one continuum material. We consider the fish fin as a three-compartment system: external fluid, apical epithelial cells, and interstitial fluid. Furthermore, owing to the geometry of the fin and for simplicity, we describe the larval fish fin as a one-dimensional system oriented along $x$ axis with a length $L$.

### 2. Ion transport and diffusion

Ions are transported between the three different compartments: we denote by $j_E^{\pm}(x)$ the ion flux from the exterior into the apical surface of the epithelium, by $j_I^{\pm}(x)$ the flux from the basal surface of the epithelium to the interstitial fluid, and by $j_p^{\pm}(x)$ the flux from the exterior to the interstitial fluid through tight junctions. The superscript $\pm$ refers to $Na^+$ and $Cl^-$, respectively. These ion fluxes read:

$$j_E^{\pm} = \beta\lambda_E^{\pm}\left(\mu_E^{\pm} - \mu_c^{\pm}\right) - S_E^{\pm}, \tag{1}$$

$$j_I^{\pm} = \beta\lambda_I^{\pm}\left(\mu_c^{\pm} - \mu_I^{\pm}\right) + S_I^{\pm}, \tag{2}$$

$$j_p^{\pm} = \beta\lambda_p^{\pm}\left(\mu_E^{\pm} - \mu_I^{\pm}\right), \tag{3}$$

where $\mu^{\pm} = k_B T \log \rho^{\pm} \pm q\Phi$ is the electrochemical potential of the ions with concentration $\rho^{\pm}$ and electric potential $\Phi$ [32]. We have also introduced the temperature $T$, the Boltzmann constant $k_B$, $\beta = 1/k_B T$, and the electric charge of an electron $q$. In Eqs. (1)-(3), we denote by $S_{E,I}^{\pm}$ the ion pump rates. And the rates of leakage through ion channels are $\lambda_{E,I}^{\pm}$, and the leak through tight junctions is $\lambda_p^{\pm}$.

Within the apical cell layer, ions diffuse between cells through gap junctions. At a coarse-grained level and averaging over the fin cross-section, it reads:

$$\partial_t \rho_c^{\pm} = D_c^{\pm}\partial_x^2 \rho_c^{\pm} \pm q\beta D_c^{\pm}\partial_x\left[\rho_c^{\pm}\partial_x \Phi_c\right] + \frac{1}{l_c}(j_E^{\pm} - j_I^{\pm}), \tag{4}$$

where $\rho_c^{\pm}(x,t)$ are the positive and negative ion concentrations in the apical cells averaged over the cross-section, $D_c^{\pm}$ is the effective diffusion constant of ions in the epithelium through gap junctions and $\Phi_c(x,t)$ is the electric potential of the cells with respect to the external medium. In addition, the source term in Eq. (4) accounts for the ion transport discussed in Eqs. (1)-(3), and $l_c$ is the area to perimeter ratio of the apical cell layer cross-section.

Similarly, the ion dynamics in the interstitial fluid is given by

$$\partial_t \rho_I^{\pm} = D_I^{\pm}\partial_x^2 \rho_I^{\pm}(x) \pm q\beta D_I^{\pm}\partial_x\left[\rho_I^{\pm}\partial_x \Phi_I\right] + \frac{1}{l_I}(j_p^{\pm} + j_I^{\pm}), \tag{5}$$

where $\rho_I^{\pm}(x,t)$ are the positive and negative ion concentrations in the interstitial fluid averaged over the cross-section, $D_I^{\pm}$ is the diffusion constant of the ion in the interstitial fluid, $\Phi_I(x,t)$ is the electric potential in the interstitial fluid with respect to the external medium and $l_I$ is the area to perimeter ratio of the internal cross-section.

### II. MULTI-TIMESCALE ELECTRICAL RESPONSE TO INJURY

We now discuss the electric response following an injury, focusing on the dynamics in the interstitial fluid. We consider that a cut of the fin is performed at time $t = 0$ at position $x = 0$, with the external medium at $x < 0$ and the interstitial fluid in the region $x > 0$.

#### 1. Electro-diffusive dynamics in the interstitial fluid

The epithelium acts as a capacitor. Ions pumps build an electric potential difference across the epithelium that leads to a net charge per unit area $\sigma_I$ on the epithelial surface facing the interstitial fluid, which reads:

$$\sigma_I = C\Phi_I, \tag{6}$$

where $C$ is the capacitance of the epithelium per unit area [33]. In addition, integrating the ion concentration along the fin cross-section, and equating it to the total charge difference across the epithelium, we obtain:

$$Aq(\rho_I^+ - \rho_I^-) = \sigma_I L_p, \tag{7}$$

where $A$ is the cross-section area and $L_p$ its perimeter. Defining the concentration difference $\rho_d \equiv \rho_I^+ - \rho_I^-$ and the dimensionless electric potential $\phi_I \equiv q\beta\Phi_I$ Eq. 7 can be written as:

$$\rho_d = c_0\phi_I, \tag{8}$$

where $c_0 = L_p C/A\beta q^2$. It can be rewritten as $c_0 = n_0/l_I$, with $qn_0 = C/\beta q$ the surface charge density that leads to a potential difference of $k_B T/q \simeq 25\,\text{mV}$, and $l_I = A/L_p$ the area to perimeter ratio of the fin cross-section. For $C \approx 0.01F/m^2$ and $l_I = 10\mu m$ we get $c_0 \simeq 35\text{nM}$.

We can now obtain the dynamics of the total ion concentration $\rho_s \equiv \rho_I^+ + \rho_I^-$ and of the dimensionless electric potential $\phi_I$ in the bulk of the interstitial fluid. For this purpose, we add and substract the dynamics of positive and negative ions in Eq. (5) to obtain:

$$\partial_t\rho_s = D\partial_{xx}\rho_s + d\partial_{xx}\rho_d + d\partial_x(\rho_s\partial_x\phi_I) + D\partial_x(\rho_d\partial_x\phi_I) + \frac{1}{l_I}\left(j^+ + j^-\right), \tag{9a}$$

$$\partial_t\rho_d = D\partial_{xx}\rho_d + d\partial_{xx}\rho_s + d\partial_x(\rho_d\partial_x\phi_I) + D\partial_x(\rho_s\partial_x\phi_I) + \frac{1}{l_I}\left(j^+ - j^-\right), \tag{9b}$$

where we have defined $D \equiv (D^+ + D^-)/2$, $d = (D^+ - D^-)/2$ and $j^\pm = j_I^\pm + j_p^\pm$. Substituting Eq. (8) in the above equations we get:

$$\partial_t \rho_s = D\partial_{xx}\rho_s + dc_0\partial_{xx}\phi_I + d\partial_x(\rho_s\partial_x\phi_I) + Dc_0\partial_x(\phi_I\partial_x\phi_I) + \frac{1}{l_I}\left(j^+ + j^-\right), \tag{10a}$$

$$c_0\partial_t\phi_I = Dc_0\partial_{xx}\phi_I + d\partial_{xx}\rho_s + dc_0\partial_x(\phi_I\partial_x\phi_I) + D\partial_x(\rho_s\partial_x\phi_I) + \frac{1}{l_I}\left(j^+ - j^-\right), \tag{10b}$$

The ion fluxes $j^\pm_{E,I,p}$ can be linearized around the steady state prior to the cut. For this purpose, we write $\rho_I^\pm = \rho_{I0}^\pm + \delta\rho_I^\pm$, $\rho_c^\pm = \rho_{c0}^\pm + \delta\rho_c^\pm$, $\phi_I = \phi_{I0} + \delta\phi_I$ and $\phi_c = \phi_{c0} + \delta\phi_c$, while we consider that $\rho_E^\pm$ remains fixed. At steady state prior to the cut, the total fluxes $j_0^\pm$ vanish. At linear order in the perturbation, the total fluxes after the cut thus read:

$$j^\pm = 0 + \delta j^\pm = \lambda_I^\pm\left(\frac{\delta\rho_c^\pm}{\rho_{c0}^\pm} - \frac{\delta\rho_I^\pm}{\rho_{I0}^\pm} \pm \delta\phi_c \mp \delta\phi_I\right) + \lambda_p^\pm\left(-\frac{\delta\rho_I^\pm}{\rho_{I0}^\pm} \mp \delta\phi_I\right). \tag{11}$$

Using electroneutrality condition in the bulk of the fluid and of the cell, we have $\rho_{I0}^+ \simeq \rho_{I0}^-$ and $\rho_{c0}^+ \simeq \rho_{c0}^-$. In addition, in the limit of high permeability of tight junction in comparison to the permeabiltiy of the cell ($\lambda_p^\pm \gg \lambda_I^\pm$), the source terms in Eq. (10) can be written as:

$$(j^+ + j_-)/l_I = -\Lambda_s\delta\rho_s/\rho_{I0} - \lambda_d\delta\phi_I\,, \tag{12a}$$

$$(j^+ - j_-)/l_I = -\lambda_d\delta\rho_s/\rho_{I0} - \Lambda_s\delta\phi_I\,, \tag{12b}$$

where we have defined $\delta\rho_s = \delta\rho_I^+ + \delta\rho_I^-$, $\rho_{I0} = \rho_{I0}^+ + \rho_{I0}^- \simeq 2\rho_{I0}^+$, $\Lambda_s = (\lambda_p^+ + \lambda_p^-)/l_I$, and $\lambda_d = (\lambda_p^+ - \lambda_p^-)/l_I$. Finally, injecting Eq. (12) into Eq. (10), dividing by $\rho_{I0}$, linearizing the nonlinear electrodiffusive terms and using the fact that $\rho_{I0} \gg c_0$, we obtain:

$$\partial_t c = D\partial_{xx}c + d\partial_{xx}\phi_I - \Lambda\delta c - \lambda\delta\phi_I, \tag{13a}$$

$$r\partial_t\phi_I = d\partial_{xx}\rho_s + D\partial_{xx}\phi_I - \lambda\delta c - \Lambda\delta\phi_I, \tag{13b}$$

where we have introduced $c = \rho_s/\rho_{I0}$, $\Lambda = \Lambda_s/\rho_{I0}$, $\lambda = \lambda_d/\rho_{I0}$ and $r = c_0/\rho_{I0}$. Equations (13) are the equations given in the main text (Fig. 2b) with with $\delta c = c - 1$ and $\delta\phi_I = \phi_I - \phi_{I0}$ and using the notation $\phi_I = \Phi$, $\phi_{I0} = \Phi_0^{\text{in}}$ and $c_0^{\text{in}} = 1$.

Note that we have kept the term $r\partial_t\phi_I$ while discarding other terms proportional to $r$. This is to illustrate the clear separation of timescale in the dynamics: using $\rho_{I0} \simeq 270$ mM and the value of $c_0 \simeq 35$ nM estimated above, we obtain $r \simeq 10^{-7}$. As we discuss further in the following, the electric potential thus reaches a quasi-steady state almost instantaneously after the cut.

#### 2. Electro-diffusive dynamics in the epithelium

So far we have looked at the dynamics of ions and electric potential in the interstitial fluid. We now discuss the dynamics in the epithelium, which is closely coupled to the dynamics in the interstitial fluid. At the timescale of experiments, the transport through gap junctions does not seem to be important (see main text for discussion). Ignoring electrodiffusion through gap junctions and linearising the fluxes $j_E^{\pm}$ and $j_I^{\pm}$ in Eq. (4) as we did in the previous section, we get

$$\partial_t \rho_c^{\pm} = -\frac{(\lambda_I^{\pm} + \lambda_E^{\pm})}{l_c} \left( \frac{\delta \rho_c^{\pm}}{\rho_{c0}^{\pm}} \pm \delta \phi_c \right) + \frac{\lambda_I^{\pm}}{l_c} \left( \frac{\delta \rho_I^{\pm}}{\rho_{I0}^{\pm}} \pm \delta \phi_I \right). \quad (14)$$

By adding the positive and negative ion dynamics in cell from the above equation, taking the limit $\lambda_I^{\pm} \gg \lambda_E^{\pm}$ and using electroneutrality $\rho_c^+ \simeq \rho_c^-$, $\rho_I^+ \simeq \rho_I^-$, we get the dynamics of total ion concentration in cells as

$$\partial_t \rho_{cs} = -\Lambda_c \frac{\delta \rho_{cs}}{\rho_{c0}} - \lambda_c \delta \phi_c + \left( \Lambda_c \frac{\delta \rho_s}{\rho_{I0}} + \lambda_c \delta \phi_I \right), \quad (15)$$

where $\rho_{cs} = \rho_c^+ + \rho_c^-$, $\rho_{c0} = \rho_{c0}^+ + \rho_{c0}^- \simeq 2\rho_{c0}^+$, $\rho_{I0} = \rho_{I0}^+ + \rho_{I0}^- \simeq 2\rho_{I0}^+$, $\Lambda_c = (\lambda_I^+ + \lambda_I^-)/\rho_{I0} l_c$, and $\lambda_c = (\lambda_I^+ - \lambda_I^-)/\rho_{I0} l_c$. Similarly, by subtracting the positive and negative ion dynamics of Eq. (14) and using electroneutrality, we obtain an equation for the electric potential inside the cell w.r.t external medium, that reads:

$$\delta \phi_c = \delta \phi_I + \frac{\lambda_c}{\Lambda_c} \left( \frac{\delta \rho_s}{\rho_{I0}} - \frac{\delta \rho_{cs}}{\rho_{c0}} \right). \quad (16)$$

We see that the membrane potential depends both on the electric potential in the interstitial fluid and the concentration of ions in the cell as well as in the interstitial fluid.

### III. COMPARISON OF THE THEORY WITH THE EXPERIMENT

#### 1. Fast changes in electric potential

Since the electric potential dynamics in Eq. (13b) is million times faster than the concentration dynamics (seconds) in Eq. (13a), over timescale of 100 ms the ion concentration can be taken to be a constant and the electric potential has reached a quasi-steadystate that is given by

$$D\partial_x^2 \phi_I - \Lambda \phi_I \simeq 0. \quad (17)$$

The solution reads: $\phi_I = \phi_{ss}\left(1 - e^{-\sqrt{\Lambda/D}\,x}\right)$. Similarly, over this fast timescale of $100ms$, the concentration of ions in the cells is constant, and we see thatthe cell potential and the interstitial fluid potential are equal in this limit:

$$\phi_c \simeq \phi_I. \tag{18}$$

Therefore the membrane potential will also exhibit an exponential profile that decays over the lengthscale $l = \sqrt{D/\Lambda}$. Fitting Eq. (18) to the experimental data yields comparable characteristic lengths for both fin layers (apical: $186 \pm 39\ \mu m$, basal: $197 \pm 22\ \mu m$, Fig. 1J). Using the diffusion constants of sodium ions $D_{\mathrm{Na}^+} = 1334\ \mu\mathrm{m}^2/\mathrm{s}$ and chloride ions $D_{\mathrm{Cl}^-} = 2032\ \mu\mathrm{m}^2/\mathrm{s}$ we obtain $D = 1683\ \mu\mathrm{m}^2/\mathrm{s}$, which gives $\Lambda = 0.0486(1 \pm 0.42)\ \mathrm{s}^{-1}$ for apical cells. We can obtain the epithelium resistance $R$ using the relation: $R = k_B T/(q^2 \rho_{I0} l_I \Lambda)$ [32]. Using $\rho_{I0} = 270$ mM and $l_I = 1\ \mu$m (see Fig. S1E) and assuming similar values for the conductivity of the positive and negative ions ($\lambda_p^+ \simeq \lambda_p^-$) we estimate the epithelial conductivity of be in the range $R \simeq 200(1 \pm 0.42)\ \Omega.\mathrm{cm}^2$. This is well within the range of reported epithelial resistances [32].

#### 2. Cellular calcium response

We model the single cell calcium response as an excitable system. To capture both the fast calcium increase and its relaxation to steady-state, we considered a minimal nonlinear model for the cytosolic calcium concentration [Ca] (we omit the time and space dependence of [Ca], $s$ and $y$ for clarity):

$$\partial_t[\mathrm{Ca}] = a\Theta_{s^*}(s)[\mathrm{Ca}]\left(1 - [\mathrm{Ca}]\right) - y[\mathrm{Ca}]\ , \tag{19a}$$

$$\tau\partial_t y = [\mathrm{Ca}] - [\mathrm{Ca}]_0\,, \tag{19b}$$

where $a$ is a constant, $s(t)$ is the dynamical signal that triggers the response and $\Theta_{s^*}(s)$ is a threshold (sigmoid) function. In the following, we use a Hill activation function $\Theta_{s^*}(s) = 1/[1 + (s^*/s)^n]$ with $n \geq 1$. Note that the precise form of the threshold function is unimportant to capture the wave propagation, provided it is almost vanishing for $s < s^*$ and switches rapidly to a value close to 1 for $s > s^*$.

The auxiliary variable $y(t)$ accounts for a secondary process (calcium reabsorbtion for instance) that allows the system to relax with timescale $\tau$ to a steady state with $[\mathrm{Ca}] = [\mathrm{Ca}]_0$ and $y = 0$.

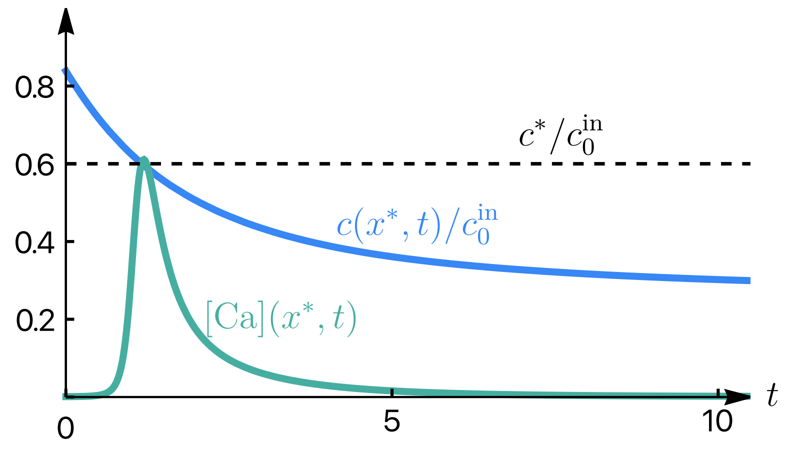


FIG. S14: Normalized ion concentration as a function of time at position $x^*$ (solid blue curve) and calcium response as a function of time at the same position $x^*$ (solid green curve). The dashline indicates the threshold value $c^*/c_0^{\rm in}$ of the nonlinear excitatory model. Values of the parameters are given in Sec. VI.

The dynamical signal $s(t)$ that triggers the response could in principle be a function of the ion concentrations and the electric potential inside the cell, which can be deduced from the ion transport model for the cells. For simplicity, and since a *diffusive* calcium front propagation is observed experimentally, we consider that the dynamical signal is proportional to the ion concentration inside the fish at cell position $x^*$. In the following, we thus take $s(t) = c(x^*, t)$. In Fig. S14, we show an example of the ionic concentration $c(x^*, t)$ and calcium response $[\rm Ca](x^*, t)$ at a fixed position $x^* = 50\ \mu$m away from the wounding site. It shows that a calcium response is triggered when the concentration $c(x^*, t)$ is close to the threshold value $c(x^*, t) = c^*$.

#### 3. Calcium activation wave in the epithelium

A propagating calcium wave is obtained from our model by considering the dynamical activation due to the diffusing ions in the interstitial fluid. For comparison with the experimental data, we consider $N$ cells indexed by $i$, and located at positions $x_i$ from the wound. The ion dynamics at these positions (obtained by solving Eq. (13)) triggers a calcium response (obtained by solving Eq. (19)) which is displayed in Fig. 2C (top panel) of the main text. From those calcium profiles, we then obtain the calcium front position as a function of time. The front position is defined as the position $x^*(t)$ at which the calcium concentration reaches an arbitrary threshold value[1] $[\mathrm{Ca}](x^*, t) = [\mathrm{Ca}]_{\mathrm{tr}}$, with $[\mathrm{Ca}]_{\mathrm{tr}} = 0.05$ in the following. The front position displays a diffusive behavior, see Fig. 2D of the main text. The apparent diffusion constant of the propagating calcium front depends on the threshold value $s^*$ in Eq. (19). We discuss this property in details in Sec. V, and show that this feature stems from the diffusing nature of the underlying signal and not from the model we considered. Interestingly, cell-level variability of this threshold may explain the spreading of the diffusive calcium signal observed experimentally. We discuss this point in Sec. III 4.

To calculate the ion concentration profile $c(x, t)$, for simplicity, we use $d = 0$ and $\lambda = 0$ in the electro-diffusion equations (Eq. (13)) and took $D = 1845$ $\mu$m$^2$/s and $\Lambda = D/l^2 \simeq 0.532$ s$^{-1}$ (with $l = 186$ $\mu$m) according to the measured values in the experiments.

The average shape of the calcium activation profiles for each cell row at a distance $x$ from the wound, obtained using the ion concentration profile and calcium dynamics given by Eq. (19), matches well with the activation profiles from the experiment but does not capture accurately the relaxation of the calcium signal (see Fig. 2C). Better agreement with the experimental calcium response in cells as a function of cell position $x$ is obtained with a rescaled calcium signal $\Delta F(x, t) = \Delta[\mathrm{Ca}](x, t)(1 - x/l)$ (see Fig. S15). The match between theory and experiment can be further improved if we allow the parameters $a$, $\tau$ to vary as a function of distance $x$.

#### 4. Cell-cell variability of the calcium activation threshold can explain noisy calcium activation wave

To discuss cell-cell variability at the light of the model, we considered $N$ cells located at positions $x_\alpha$ along the fin. Each of those cells may have different parameters $a_\alpha$, $\tau_\alpha$, $n_\alpha$ and $s^*_\alpha$ for the nonlinear calcium dynamics given by Eq. (19). We consider the case where $s^*_\alpha$ is non-homogeneous, while parameters $a$, $\tau$, $n$ are homogeneous for all cells. To model the cell-to-cell variability of activation threshold, for each cell, we draw $s^*$ from a Gaussian distribution with mean $\langle s^* \rangle = s^*_0$ and variance $\sigma$.

The resulting noisy propagation of the calcium activation front is shown in Fig. S16 for $N = 1000$, $s^*_0 = 0.6c^{\mathrm{in}}_0$ and $\sigma = 0.1c^{\mathrm{in}}_0$. The qualitative similarity of this plot to the

[1] Changing the threshold value $[\mathrm{Ca}]_{\mathrm{tr}}$ does not change significantly the diffusive behaviour that we observe. Setting a larger (respectively smaller) values of this threshold will however decrease (resp. increase) the length of propagation of this front.

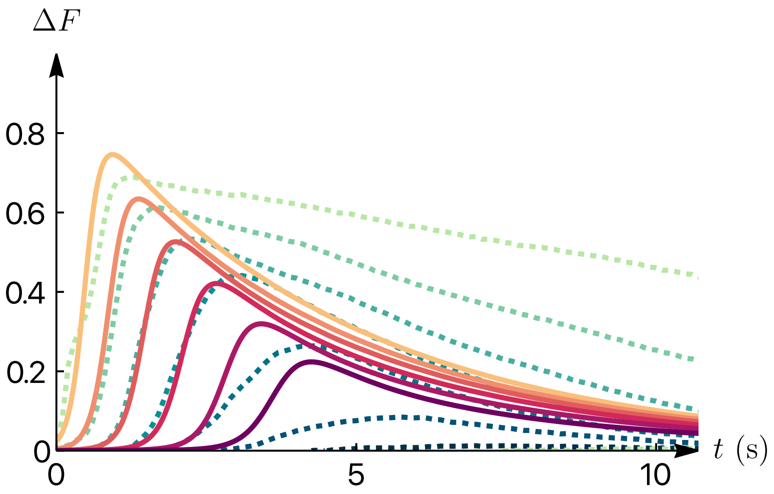


FIG. S15: Cytosolic $Ca^{2+}$ intensity as a function of time at different positions away from the wound from the nonlinear model of cellular calcium response (thick lines in shades of orange), and from the experiments (dashed lines in shades of green). The experimental data is the same as the one displayed in Fig. 2C (bottom) of the main text.

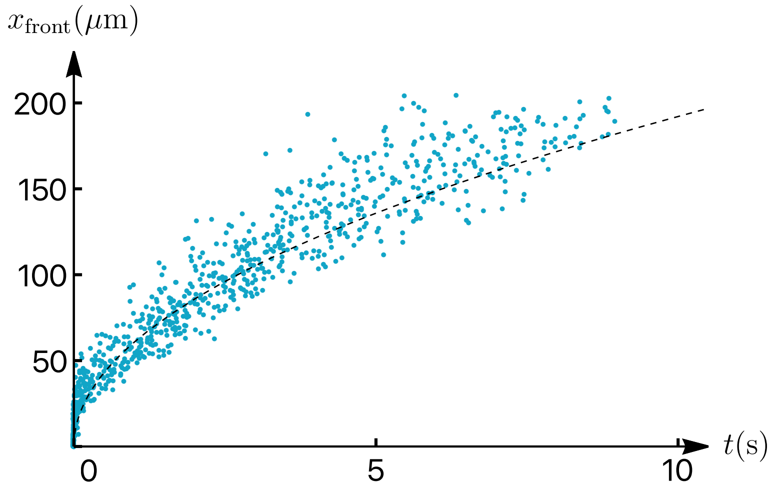


FIG. S16: Noisy propagation of the calcium wave front. Blue dots indicate the position $x_{\text{front}}$ where a calcium response has been activated as a function of time $t$ obtained using the model described in Sec. III 4. The black dashed line shows $\sqrt{2Dt}$.

experimental data presented in Fig. 1F suggests that the noisy calcium activation front originates from cell-to-cell variability in the threshold parameter $s^*$.

### IV. EFFECT OF ADDING POTASSIUM CHLORIDE TO THE EXTERNAL MEDIUM ON THE ION ELECTRO-DIFFUSIVE DYNAMICS

#### 1. Effect of potassium chloride supplement on epithelial electric potential

To understand the effect of adding KCl on the membrane potential of the epithelial cells we can extend the framework developed in Sec. I to include the dynamics of $K^+$ along with $Na^+$ and $Cl^-$ considered before. Following Eq. (4), the dynamics of the ions inside the epithelium is given by

$$\partial_t \rho_c^i = D_c^i \partial_x^2 \rho_c^i + z_i q\beta D_c^i \partial_x \left[\rho_c^i \partial_x \Phi_c\right] + \frac{1}{l_c}(j_E^i - j_I^i), \tag{20}$$

where $i \in \{\mathrm{Na}^+, \mathrm{K}^+, \mathrm{Cl}^-\}$ is the label for the ion type and $\mathbf{z} = (1, 1, -1)$ is the valency of the ions. Since KCl is homogeneously distributed in the external medium, we do not expect a spatial variation in the ion concentration in the epithelium; therefore, we can drop the gradient terms in Eq. (20). The conservation of electric current implies $\partial_t \rho_c^{Na^+} + \partial_t \rho_c^{K^+} = \partial_t \rho_c^{Cl^-}$. Substituting the dynamics of the three ions as given by Eq. (20) in this we get:

$$j_E^{Na^+} - j_I^{Na^+} + j_E^{K^+} - j_I^{K^+} = j_E^{Cl^-} - j_I^{Cl^-}. \tag{21}$$

The ion fluxes are given by: $j_E^i = \beta\lambda_E^i\left(\mu_E^i - \mu_c^i\right) - S_E^i$ and $j_I^i = \beta\lambda_I^i\left(\mu_c^i - \mu_I^i\right) + S_I^i$, where the notations are the same as that in Eq. (1). Substituting these fluxes in Eq. (21), we get

$$q\beta\Phi_c = \frac{1}{\sum_i\left(\lambda_E^i + \lambda_I^i\right)}\sum_i\left(\lambda_I^i q\beta\Phi_I + \lambda_E^i z_i \log\frac{\rho_E^i}{\rho_c^i} + \lambda_I^i z_i \log\frac{\rho_I^i}{\rho_c^i} - z_i S_E^i - z_i S_I^i\right). \tag{22}$$

Similarly, following Eq. (5), the dynamics of the ions in the interstitial fluid is given by:

$$\partial_t \rho_I^i = D_I^i \partial_x^2 \rho_I^i(x) + z_i q\beta D_I^i \partial_x \left[\rho_I^i \partial_x \Phi_I\right] + \frac{1}{l_I}(j_p^i + j_I^i). \tag{23}$$

For a homogeneous system, substituting the ion dynamics from Eq. (23) in the current conservation condition $\partial_t \rho_I^{Na^+} + \partial_t \rho_I^{K^+} = \partial_t \rho_I^{Cl^-}$, we get:

$$q\beta\Phi_I = \frac{1}{\sum_i\left(\lambda_I^i + \lambda_p^i\right)}\sum_i\left(\lambda_I^i q\beta\Phi_c + \lambda_I^i z_i \log\frac{\rho_c^i}{\rho_I^i} + \lambda_p^i z_i \log\frac{\rho_E^i}{\rho_I^i} + z_i S_I^i\right). \tag{24}$$

To understand the immediate response of adding KCl to the external medium to $\Phi_c$ and $\Phi_I$ we fix the concentration of all ions except the extracellular $K^+$ and $Cl^-$ and simultaneously solve the two linear equations Eq. (22) and Eq. (24) to obtain the $\Phi_c$ and $\Phi_I$. The concentration of $K^+$ and $Cl^-$ is negligible compared to the perturbation. The change in the potential when KCl of concentration of $\delta\rho$ is added to the external medium while keeping all other concentration constant reads:

$$q\beta\delta\Phi_c = \frac{1}{(\lambda_E + \lambda_I)}\left(\lambda_I q\beta\delta\Phi_I + \left(\lambda_E^{K^+} - \lambda_E^{Cl^-}\right)\log\delta\rho\right) \tag{25}$$

$$q\beta\delta\Phi_I = \frac{1}{(\lambda_I + \lambda_p)}\left(\lambda_I q\beta\delta\Phi_c + \left(\lambda_p^{K^+} - \lambda_p^{Cl^-}\right)\log\delta\rho\right), \tag{26}$$

where $\lambda_{E,I,p} = \sum_i \lambda_{E,I,p}^i$. Solving the two equations we get the change in epithelia potential w.r.t external medium as

$$q\beta\delta\Phi_c = \frac{\left(\left(\lambda_p^{K^+} - \lambda_p^{Cl^-}\right)\lambda_I + \left(\lambda_E^{K^+} - \lambda_E^{Cl^-}\right)(\lambda_p + \lambda_I)\right)\log\delta\rho}{\lambda_E\lambda_I + \lambda_p\lambda_I + \lambda_E\lambda_p}. \tag{27}$$

The sign of the membrane potential change $\delta\Phi_c$ depends on the relative permeabilities. If $\left(\lambda_p^{K^+} - \lambda_p^{Cl^-}\right)\lambda_I < \left(\lambda_E^{K^+} - \lambda_E^{Cl^-}\right)(\lambda_p + \lambda_I)$, the epithelium depolarizes upon addition of KCl in the external medium. This calculation shows that adding KCl to the system modulates the membrane potential. This is consistent with our observation that adding KCl leads to changes in proliferation regulated via the Voltage Sensing Phosphatase VSP (Figure 4F-K).

#### 2. Slower diffusion of sodium ion in the presence of potassium ion

To understand the diffusive dynamics of ions in the potential supplemented case we analyze a one dimension electro-diffusion model with three ions: $Na^+$, $K^+$, and $Cl^-$ in a channel. The dynamics of the ions is given by the following equations.

$$\partial_t\rho_{Na} = \nabla\cdot\left[D_{Na}\rho_{Na}\nabla\left(\log\rho_{Na} + q\beta\Phi\right)\right], \tag{28}$$

$$\partial_t\rho_K = \nabla\cdot\left[D_K\rho_K\nabla\left(\log\rho_K + q\beta\Phi\right)\right], \tag{29}$$

$$\partial_t\rho_{Cl} = \nabla\cdot\left[D_{Cl}\rho_{Cl}\nabla\left(\log\rho_{Cl} - q\beta\Phi\right)\right], \tag{30}$$

where $\rho_{Na}, \rho_K, \text{and}, \rho_{Cl}$ are the concentration of $Na^+$, $K^+$, and $Cl^-$, respectively. At the timescale and lengthscale of interest we can take the medium to be electroneutral, which imposes the constraint $\rho_{Na} + \rho_K = \rho_{Cl}$ and $\partial_t\rho_{Na} + \partial_t\rho_K = \partial_t\rho_{Cl}$. Substituting the ion dynamics in this last equation, we get the following expression for the electric potential as a function of sodium and potassium ion concentrations:

$$\nabla\left(\left(D_{Na}\rho_{Na} + D_K\rho_K + D_{Cl}\rho_{Cl}\right)\nabla(q\beta\Phi) + \nabla\left(D_{Na}\rho_{Na} + D_K\rho_K - D_{Cl}\rho_{Cl}\right)\right) = 0. \tag{31}$$

Integrating Eq. (31) and setting the flux to be zero at one of the far end we get:

$$\left(\bar{D}_{Na}\rho_{Na} + \bar{D}_K\rho_K\right)\nabla(q\beta\Phi) = -\tilde{D}_{Na}\rho_{Na}\nabla\log\rho_{Na} - \tilde{D}_K\rho_K\nabla\log\rho_K, \tag{32}$$

where we have used the electroneutrality condition: $\rho_{Na} + \rho_K = \rho_{Cl}$, and defined $\bar{D}_{Na} \equiv D_{Na}+D_{Cl}$, $\bar{D}_K \equiv D_K+D_{Cl}$, $\tilde{D}_{Na} \equiv D_{Na}-D_{Cl}$, and $\tilde{D}_K \equiv D_K-D_{Cl}$. We can now substitute the electric field from Eq. (32) in Eqs. (28) and (29) to obtain an effective dynamics for the sodium and potassium ions that reads:

$$\partial_t\rho_{Na} = \nabla\cdot\frac{D_{Na}\left(2D_{Cl}\rho_{Na} + \bar{D}_K\rho_K\right)\nabla\rho_{Na} - D_{Na}\tilde{D}_K\rho_{Na}\nabla\rho_K}{\left(\bar{D}_{Na}\rho_{Na} + \bar{D}_K\rho_K\right)}, \tag{33}$$

$$\partial_t\rho_K = \nabla\cdot\frac{-D_K\tilde{D}_{Na}\rho_K\nabla\rho_{Na} + D_K\left(2D_{Cl}\rho_K + \bar{D}_{Na}\rho_{Na}\right)\nabla\rho_K}{\left(\bar{D}_{Na}\rho_{Na} + \bar{D}_K\rho_K\right)}. \tag{34}$$

Since the diffusion constants of potassium and chloride are very similar, we take them to be equal ($\tilde{D}_K \simeq 0$) for simplicity. In addition, linearizing around the state $\rho_{Na} = \rho^0_{Na} + \delta\rho_{Na}$ and $\rho_K = \rho^0_K + \delta\rho_K$, and considering $\rho^0_{Na} \gg \rho^0_K$, the fluxes simplify to

$$\partial_t\rho_{Na} = \frac{2D_{Na}D_{Cl}}{\bar{D}_{Na}}\left[1 + \left(\frac{\bar{D}_K}{2D_{Cl}}\frac{\tilde{D}_{Na}}{\bar{D}_{Na}}\right)\frac{\rho^0_K}{\rho^0_{Na}}\right]\nabla^2\rho_{Na}, \tag{35}$$

$$\partial_t\rho_K = D_K\nabla^2\rho_K - \frac{D_K\tilde{D}_{Na}\rho^0_K}{\bar{D}_{Na}\rho^0_{Na}}\nabla^2\rho_{Na}. \tag{36}$$

The effective diffusion of sodium ion in this limit is

$$D^{\text{eff}}_{Na} = \frac{2D_{Na}D_{Cl}}{\bar{D}_{Na}}\left[1 + \left(\frac{\bar{D}_K}{2D_{Cl}}\frac{\tilde{D}_{Na}}{\bar{D}_{Na}}\right)\frac{\rho^0_K}{\rho^0_{Na}}\right]. \tag{37}$$

This demonstrates that adding extra KCl in the extracellular medium alters the diffusion coefficient of the ions in the IF. This is consistent with the observation that the $Ca^{2+}$ diffusive wavefront is altered upon addition of extraembryonic KCl. Since $\tilde{D}_{Na} < 0$, we see that the effective diffusion constant of sodium is reduced in the presence of potassium ions.

In our model with one cation and one anion, total ion concentration and cation concentration are proportional. Thus, calcium dynamics driven by $s(x,t) = c(x,t)$ or $s(x,t) = c(x,t)/2$ yield similar responses. In contrast, in the three-ion model different ions acting as driving signal $s(x,t)$ may give different effective diffusion constant of calcium activation wavefront. If calcium release is primarily triggered by $Na^+$, adding KCl reduces the $Na^+$ diffusivity, thus slowing calcium activation wave propagation. Experimental observations of slower calcium diffusion supports this idea (Fig. 2H). Moreover, the threshold $s^*$, may also be altered by the change in external ionic environment, leading to change in the effective diffusion constant.

### V. DIFFUSION FRONT AND EFFECTIVE DIFFUSION CONSTANT

In this section we discuss the effective diffusion constant of a propagating front. In a first subsection, we show that such a dependency is generic for a diffusive system. In a second subsection, we discuss this dependency for the calcium wave model of Sec. III.

#### 1. Simple diffusion with threshold

We discuss the dynamics of the position of a diffusive front, that is, the dynamics of the position $x^*$ at which a given value $c^*$ of the concentration is reached. We consider an "infinite fish" geometry: The fish is located in the $x \leq 0$ half plane with initial inner concentration $c_0$, and a wound is located at position $x = 0$. The outer medium ($x > 0$) is assumed to have a vanishing concentration. We consider simple diffusion $\partial_t c = D\partial_x^2 c$ with initial condition $c(x, t = 0) = c_0\Theta(-x)$, and boundary condition $c(x = -\infty, t) = c_0$ and $c(x = +\infty, t) = 0$. The simple geometry allows for an analytical solution:

$$c(x,t) = \frac{c_0}{2}\left[1 - \mathrm{erf}(x/2\sqrt{Dt})\right] , \tag{38}$$

with the error function $\mathrm{erf}(x) = 2\int_0^x \mathrm{d}u\, \mathrm{e}^{-u^2}/\sqrt{\pi}$.

The front position $x^*$ has a diffusive dynamics with an effective diffusion constant $x^*(t) = \sqrt{2D_{\mathrm{eff}}t}$. This effective constant can be obtained directly by imposing $c(x^*, t) = c^*$ and yields $D_{\mathrm{eff}} = 2D\left[\mathrm{erf}^{-1}(1 - 2c^*/c_0)\right]^2$ where $\mathrm{erf}^{-1}$ is the inverse of the erf function.

Importantly, this shows that the effective diffusion constant depends on the value $c^*$ chosen for the threshold. We display this dependence in Fig. S17.

#### 2. Effective diffusion constant in the tissue model

The generic dependency of the effective diffusion constant discussed above also applies to our (simplified) tissue model discussed in Sec. III.

Depending on the value of the excitable model threshold $s^* = 1 - c^*$, a diffusive behavior with different effective constants can be observed (see Fig. S18). Cell-level variability of this threshold may explain the spreading of the diffusive calcium signal observed experimentally.

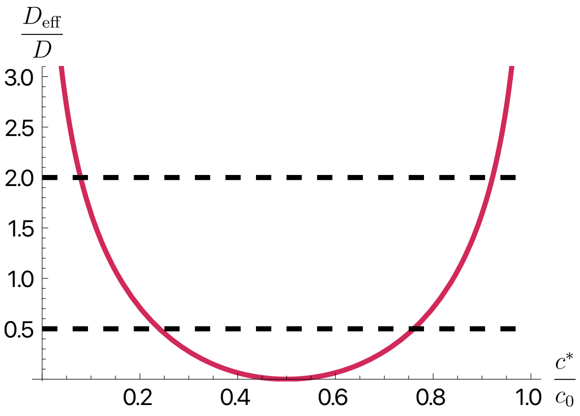


FIG. S17: Effective diffusion constant of the front position as a function the concentration threshold (solid red curve). The dashed lines indicate the range where the effective diffusion constant is between half the value of the diffusion process and twice its value.

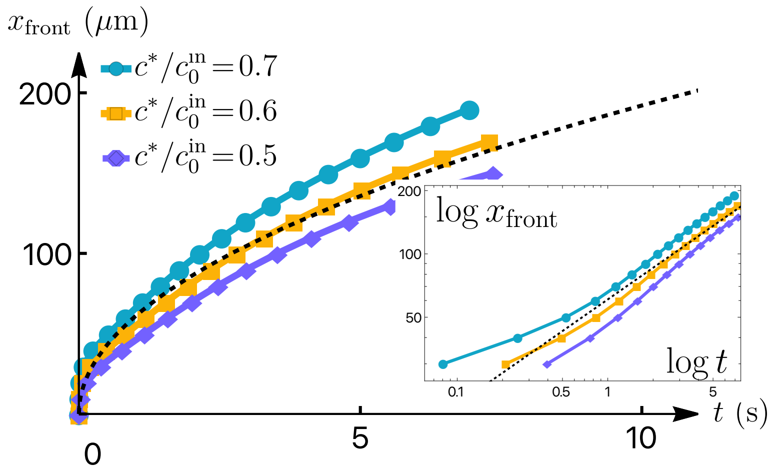


FIG. S18: Calcium wave front position $x_{\mathrm{front}}$ as a function of time $t$. Different colors indicate different values of the threshold of the cellular calcium response model Eq. (19). The black dashed line shows $\sqrt{2Dt}$.Inset shows the same information in log-log scale.

### VI. PARAMETERS USED IN THE FIGURES AND MOVIE

In Fig. 2B,C,D of the main text as well as in Figs. S14, S18, S16, S15 and Movie S4 of the SI, we used the following parameters for the ion dynamics: $D = 1845\ \mu\mathrm{m}^2/\mathrm{s}$, $\Lambda = 0.532\ \mathrm{s}^{-1}$, $c_0^{\mathrm{out}}/c_0^{\mathrm{in}} = 10/270$, $d = 0$, and $\lambda = 0$.

In all those figures and movie, we used the following parameters for the cellular calcium response model: $a = 30$, $\tau = 0.66$ s, $[\mathrm{Ca}]_0 = 10^{-3}$, $c^*/c_0^{\mathrm{in}} = 0.6$ and $n = 4$, except for Fig. S15, for which we used the following parameters: $a = 10$, $\tau = 5$ s, $[\mathrm{Ca}]_0 = 10^{-3}$,

$c^*/c_0^{\text{in}} = 0.3$ and $n = 2$.

---

[1] M. Westerfield, *The Zebrafish Book. A Guide for the Laboratory Use of Zebrafish (Danio Rerio)* (Univ. of Oregon Press, Eugene, USA, 2000).

[10] T.-W. Chen, T. J. Wardill, Y. Sun, S. R. Pulver, S. L. Renninger, A. Baohan, E. R. Schre-

iter, R. A. Kerr, M. B. Orger, V. Jayaraman, L. L. Looger, K. Svoboda, and D. S. Kim, Ultrasensitive fluorescent proteins for imaging neuronal activity, Nature **499**, 295 (2013).

[11] C. Mosimann, C. K. Kaufman, P. Li, E. K. Pugach, O. J. Tamplin, and L. I. Zon, Ubiquitous transgene expression and Cre-based recombination driven by the *ubiquitin* promoter in zebrafish, Development **138**, 169 (2011).

[12] J. M. Daane, N. Blum, J. Lanni, H. Boldt, M. K. Iovine, C. W. Higdon, S. L. Johnson, N. R. Lovejoy, and M. P. Harris, Modulation of bioelectric cues in the evolution of flying fishes, Curr. Biol. **31**, 5052 (2021).

[13] A. S. Abdelfattah, J. Zheng, A. Singh, Y.-C. Huang, D. Reep, G. Tsegaye, A. Tsang, B. J. Arthur, M. Rehorova, C. V. Olson, Y. Shuai, L. Zhang, T.-M. Fu, D. E. Milkie, M. V. Moya, T. D. Weber, A. L. Lemire, C. A. Baker, N. Falco, Q. Zheng, J. B. Grimm, M. C. Yip, D. Walpita, M. Chase, L. Campagnola, G. J. Murphy, A. M. Wong, C. R. Forest, J. Mertz, M. N. Economo, G. C. Turner, M. Koyama, B.-J. Lin, E. Betzig, O. Novak, L. D. Lavis, K. Svoboda, W. Korff, T.-W. Chen, E. R. Schreiter, J. P. Hasseman, and I. Kolb, Sensitivity optimization of a rhodopsin-based fluorescent voltage indicator, Neuron **111**, 1547 (2023).

[20] A. Kawakami, T. Fukazawa, and H. Takeda, Early fin primordia of zebrafish larvae regenerate

by a similar growth control mechanism with adult regeneration, Dev. Dyn. **231**, 693 (2004).

[21] R. Mateus, T. Pereira, S. Sousa, J. E. de Lima, S. Pascoal, L. S. de, and A. Jacinto, In Vivo Cell and Tissue Dynamics Underlying Zebrafish Fin Fold Regeneration, PLOS ONE **7**, e51766 (2012).

[22] M. Pachitariu and C. Stringer, Cellpose 2.0: How to train your own model, Nat. Methods **19**, 1634 (2022).

[23] J.-Y. Tinevez, Hungarian based particle linking (2005), matlab Central File Exchange.

[24] S. Stewart, H. K. Le Bleu, G. A. Yette, A. L. Henner, A. E. Robbins, J. A. Braunstein, and K. Stankunas, *Longfin* causes *cis* -ectopic expression of the *kcnh2a ether-a-go-go* K+ channel to autonomously prolong fin outgrowth, Development **148**, dev199384 (2021).

[25] J. M. Daane, J. Lanni, I. Rothenberg, G. Seebohm, C. W. Higdon, S. L. Johnson, and M. P. Harris, Bioelectric-calcineurin signaling module regulates allometric growth and size of the zebrafish fin, Sci. Rep. **8**, 10391 (2018).

[26] C. Yi, T. W. Spitters, E. A.-D. A. Al-Far, S. Wang, T. Xiong, S. Cai, X. Yan, K. Guan, M. Wagner, A. El-Armouche, and C. L. Antos, A calcineurin-mediated scaling mechanism that controls a K+-leak channel to regulate morphogen and growth factor transcription, eLife **10**, e60691 (2021).

[27] S. Perathoner, J. M. Daane, U. Henrion, G. Seebohm, C. W. Higdon, S. L. Johnson, C. Nüsslein-Volhard, and M. P. Harris, Bioelectric Signaling Regulates Size in Zebrafish Fins, PLoS Genet. **10**, e1004080 (2014).

[28] J. Liu, Fast calcium signals upon fin injury analysis codes (2025).

[29] J. Liu, Membrane potential reporter analysis upon fin injury (2025).

[30] J. Schindelin, I. Arganda-Carreras, E. Frise, V. Kaynig, M. Longair, T. Pietzsch, S. Preibisch, C. Rueden, S. Saalfeld, B. Schmid, J.-Y. Tinevez, D. J. White, V. Hartenstein, K. Eliceiri, P. Tomancak, and A. Cardona, Fiji: An open-source platform for biological-image analysis, Nat. Methods **9**, 676 (2012).

[31] E. Nerli, Proliferation analysis fin (2025).

[32] A. Torres-Sánchez, M. Kerr Winter, and G. Salbreux, Tissue hydraulics: Physics of lumen formation and interaction, Cells & Development **168**, 203724 (2021).

[33] Y. Mori, From Three-Dimensional Electrophysiology to the Cable Model: An Asymptotic Study, arXiv: 0901.3914 10.48550/arXiv.0901.3914 (2009), arXiv:0901.3914.
